# Anchored Phylogenomics of Angiosperms I: Assessing the Robustness of Phylogenetic Estimates

**DOI:** 10.1101/086298

**Authors:** Chris Buddenhagen, Alan R. Lemmon, Emily Moriartya Lemmon, Jeremy Bruhl, Jennifer Cappa, Wendy L. Clement, Michael J. Donoghue, Erika J. Edwards, Andrew L. Hipp, Michelle Kortyna, Nora Mitchell, Abigail Moore, Christina J. Prychid, Maria C. Segovia-Salcedo, Mark P. Simmons, Pamela S. Soltis, Stefan Wanke, Austin Mast

## Abstract

An important goal of the angiosperm systematics community has been to develop a shared approach to molecular data collection, such that phylogenomic data sets from different focal clades can be combined for meta-studies across the entire group. Although significant progress has been made through efforts such as DNA barcoding, transcriptome sequencing, and whole-plastid sequencing, the community current lacks a cost efficient methodology for collecting nuclear phylogenomic data across all angiosperms. Here, we leverage genomic resources from 43 angiosperm species to develop enrichment probes useful for collecting ~500 loci from non-model taxa across the diversity of angiosperms. By taking an anchored phylogenomics approach, in which probes are designed to represent sequence diversity across the group, we are able to efficiently target loci with sufficient phylogenetic signal to resolve deep, intermediate, and shallow angiosperm relationships. After demonstrating the utility of this resource, we present a method that generates a heat map for each node on a phylogeny that reveals the sensitivity of support for the node across analysis conditions, as well as different locus, site, and taxon schemes. Focusing on the effect of locus and site sampling, we use this approach to statistically evaluate relative support for the alternative relationships among eudicots, monocots, and magnoliids. Although the results from supermatrix and coalescent analyses are largely consistent across the tree, we find support for this deep relationship to be more sensitive to the particular choice of sites and loci when a supermatrix approach as employed. Averaged across analysis approaches and data subsampling schemes, our data support a eudicot-monocot sister relationship, which is supported by a number of recent angiosperm studies.

---

A longstanding aim of angiosperm systematists has been to develop a unified approach to data collection across the entire clade. Previous efforts have demonstrated the importance of community coordination (e.g., *rbcL* sequencing; Chase et al. 1993). As the systematics community moves into the genomics era, however, new technologies are enabling researchers to transition from targeting a small number of primarily organellar loci, to targeting hundreds or thousands of loci that are both broadly distributed across the genome and broadly homologous across deep clades (Mamanova et al. 2010; Griffin et al. 2011; Crawford et al. 2012; Cronn et al. 2012; Lemmon and Lemmon 2013; Davis et al. 2014; Wickett et al. 2014; Prum et al. 2015; Wicke and Schneeweiss 2015). Although progress has been made in plants recently using other approaches such as whole genome sequencing, whole-plastid sequencing (Ruhfel et al. 2014), and transcriptome sequencing (Burleigh et al. 2011; Lee et al. 2011; Wickett et al. 2014; Zeng et al. 2014), perhaps the most promising methodology to date is hybrid enrichment, which is rapidly resolving many previously intractable branches across the Tree of Life (Stull et al. 2013; Pyron et al. 2014; Brandley et al. 2015; Crawford et al. 2015; Eytan et al. 2015; Prum et al. 2015; Bryson et al. 2016; Meiklejohn et al. 2016; Tucker et al. 2016; Young et al. 2016). This method has the advantages of cost-effectiveness, high-throughput data production, and success with historical specimens, such as herbarium samples (Bi et al. 2013; Stull et al. 2013; Bakker 2015; Beck and Semple 2015). Hybrid enrichment is a method wherein short oligonucleotides (termed “probes” or “baits”) are designed to complement target loci across the genomes of a clade (e.g., vertebrates, angiosperms, etc.). The researcher can choose the number, length, and type of loci to be targeted. In the laboratory, probes hybridize to genomic DNA libraries, facilitating enrichment of target loci relative to the remainder of the genome. The enriched DNA libraries are then sequenced on a high-throughput sequencing platform. Because of the flexible and customizable nature of the method, hybrid enrichment has the potential to unify data collection for many different types of research questions across angiosperms.

Some versions of hybrid enrichment are generally less appropriate than others for phylogenetic studies in plants. For example, the “UCE” method, as originally defined (Faircloth et al. 2012), utilizes thousands of relatively-short ultraconserved elements scattered across the genome as target regions. The use of UCEs in plants may be limited because appear to be scarce in plants compared to other systems (6 non-repetitive, invariant UCEs in plants vs. 1120 in animals; Reneker et al. 2012). An alternative method, Anchored Hybrid Enrichment (AHE; Lemmon et al. 2012; Lemmon and Lemmon 2013), ensures high enrichment efficiency by leveraging genomic and transcriptomic resources to develop probe sets that reflect the natural sequence variation occurring across a clade. By incorporating probes reflecting this phylogenetic diversity, regions with moderate levels of sequence variation can be efficiently enriched. Following recommendations from simulation studies (Leaché and Rannala 2011), this method incorporates a moderate number (~500-1000) of moderately-conserved target loci. Thus, the phylogenetic information content of the data sets generated is high, and the reasonable sequencing effort required makes it affordable for researchers to collect data from hundreds to thousands of species across evolutionary timescales. This method has been applied to diverse taxa across the Tree of Life, leading to complete or nearly complete resolution of many clades of different taxonomic depth, including birds (Prum et al. 2015; Morales et al. 2016), snakes (Pyron et al. 2014; Ruane et al. 2015), flies (Young et al. 2016), lizards (Brandley et al. 2015; Tucker et al. 2016), teleost fish (Eytan et al. 2015), frogs (Peloso et al. 2016). Given the success of AHE in many other systems, and the success to date of hybrid enrichment efforts targeting individual angiosperm clades (Table 1; e.g., Asteraceae, Mandel et al. 2014), the time seems ripe to extend AHE to angiosperm phylogenetics though development and implementation of a general angiosperm-wide enrichment resource.

**Table 1.** Recent phylogenetic or population genetic studies that used hybrid-enrichment methods, including plastid-oriented studies. Studies targeting only nuclear regions but that utilized plastid sequences found in bycatch are denoted by an asterisk (*). # Ref. Taxa refers to the number of reference taxa included in a probe design. # Sp. Copy Assessment indicates how many species the study included in their assessment of copy number for each locus included in the probe kit developed.

| Study | Basal Group | Major Group | Family | Genus | # Ref. Taxa | # Nuclear Loci Targeted | Probes Target Plastids | # Sp. Copy Assessment |
| --- | --- | --- | --- | --- | --- | --- | --- | --- |
| Parks et al. (2012) | Gymnosperms | - | Pinaceae | *Pinus* | 2 | 0 | yes | 2 |
| Stull et al. (2013) | Angiosperms | eudicots | - | - | 22 | 0 | yes | 0 |
| Mandel et al. (2014) | Angiosperms | eudicots | Asteraceae | - | 1 | 1061 | no | 4 |
| Weitmier et al. (2014) | Angiosperms | eudicots | Asteraceae | *Asclepias* | 1 | 3385 | no* | 6 |
| de Sousa et al. (2014) | Angiosperms | eudicots | Leguminosae | *Medicago* | 1 | 319 | no | 5 |
| Grover et al. (2015) | Angiosperms | eudicots | Malvaceae | *Gossypium* | 1 | 500 | no | 1 |
| Stephens et al. (2015a) | Angiosperms | eudicots | Sarraceniaceae | *Sarracenia* | 2 | 646 | no* | 2 |
| Stephens et al. (2015b) | Angiosperms | eudicots | Asteraceae | *Helianthus* | 4 | 598 | no* | 2 |
| Schmickl et al. (2016) | Angiosperms | eudicots | Oxalidaceae | *Oxalis* | 1 | 1164 | no* | 3 |
| Heyduk et al. (2016) | Angiosperms | monocots | Arecaceae | *Sabal* | 15 | 837 | yes | 15 |
| Syring et al. (2016) | Gymnosperms | - | Pinaceae | *Pinus albicaulis* | 1 | 7849 | no | 1 |
| Sass et al. (2016) | Angiosperms | monocots | Zingiberaceae | - | 8 | 494 | yes | 8 |
| **This Study** | **Angiosperms** | **All groups** | - | - | **25** | **517** | **no\*** | **43** |

Despite tremendous efforts to reconstruct the phylogeny of angiosperms, several deep nodes continue to be debated (Soltis et al. 2011; Burleigh et al. 2011; Wickett et al. 2014; Magallón et al. 2013). One reason for this difficulty is the large diversity in angiosperms— estimates suggest approximately 400,000 extant species (~300,000 described; Pimm and Joppa 2015). A second reason is the limitation of many previous studies to a small number of plastid and nuclear ribosomal markers available at the time, which do not always contain sufficient information for resolving challenging nodes (e.g., Zimmer and Wen 2012), due to processes such as incomplete lineage sorting and hybridization. A third reason is that many previous studies have only presented results from a narrow set of data subsampling and methodological conditions (e.g., Wickett et al. 2014). An especially intractable node that may be influenced by these problems is the relationship among eudicots, monocots, and magnoliids at the base of the angiosperms. Previous work has suggested all possible alternative resolutions of these three clades depending upon type of data or taxa included and analysis method (eudicots-monocots sister: Soltis et al. 2011; eudicots-magnoliids sister: Burleigh et al. 2011, Lee et al. 2011; magnoliids-monocots sister: Wickett et al. 2014). Elucidating the true history of this group and other difficult clades will require a more comprehensive approach that accounts for effects of both taxa and data inclusion.

Systematists are increasingly becoming aware of the fact that although sufficient genomic and taxonomic samplings are necessary for accurate phylogeny estimation, proper data analysis is also critical. Accurate reconstruction requires careful navigation of a phylogenetic workflow that is becoming increasingly complex and sophisticated (reviewed by Lemmon and Lemmon 2013). For example, evolutionary processes must be modeled adequately or estimates of topology and branch lengths may be biased (Posada and Crandall 1998; Lemmon and Moriarty 2004; Brown and Lemmon 2007; Lanfear et al. 2012; Hoff et al. 2016). Second, proper orthology assessment is essential to avoid including paralogous sequences (Chen et al. 2007; Lemmon and Lemmon 2013). Third, construction of complete or nearly-complete data matrices can be important for minimizing the potential effects of missing data (Lemmon 2009; Roure et al. 2013; Hosner et al. 2015). Finally, a particularly difficult aspect of phylogenomic analysis is how to accommodate multilocus data with different underlying histories. The traditional approach has been to concatenate the loci into a single supermatrix in the hope that the signal of the true underlying species history overwhelms any discrepancies among the individual histories of the loci sampled (Kluge 1989; de Queiroz and Gatesy 2007). More recently, however, theory and simulation studies have shown that explicitly accounting for incongruent gene histories (e.g., using a coalescent model or networks) can be essential for accurate estimation of species history (Edwards 2009; Morrison 2011; Mirarab and Warnow 2015; Edwards et al. 2016). Incongruence among loci has particular bearing on angiosperm systematics because relationships among some of the key groups change depending upon whether a supermatrix or coalescent (species-tree) approach is employed (e.g., Wickett et al. 2014; Edwards et al. 2016; Sun et al. 2016), although it is possible that increased taxon sampling could mitigate these effects. What is currently lacking is a framework for systematically comparing results from different analysis conditions, while also considering the sensitivity of these results to particular combinations of taxa and loci within the data set.

Here, we develop and test an angiosperm-wide hybrid enrichment resource for general use by the angiosperm systematics community in order to facilitate unification of data collection and accelerate resolution within this clade. After identifying 499 target loci with appropriate characteristics derived from Duarte et al. (2010), we develop a robust AHE probe kit that utilizes genomic resources from 25 genomes distributed broadly across angiosperms. We demonstrate the value of this resource for use at deep, intermediate, and shallow scales by carrying out hybrid enrichment across six orders and 10 plant families, and we incorporate previously published data to estimate a phylogenetic tree of 30 of the 79 angiosperm orders (APG IV 2016). Note that our sampling is designed to provide sufficient taxonomic coverage to evaluate the effectiveness of the enrichment probes, not to produce a comprehensive phylogeny of angiosperms. Independent of the phylogenetic data collection, we illustrate the potential for collecting data from functional loci by including probes for 18 selenium-tolerance loci (Freeman et al. 2010). Finally, we present results from a sensitivity analysis that will allow for a more systematic distinction between robust relationships that do not require further study versus tenuous relationships that require a more thorough investigation. Our goals are to: (1) test the efficiency of the enrichment resource and its effectiveness in resolving phylogenies and (2) provide a novel framework for evaluating the robustness of phylogenetic estimates and testing alternative topological hypotheses. The hybrid enrichment resource we present here is broad both genomically (loci are selected from throughout the genome; Suppl. Fig. 1) and taxonomically (taxa represented are distributed across the clade) in order to provide a useful tool for widespread application in angiosperm systematics studies. A practical demonstration of the application of this methodology to resolving difficult nodes below the family level is provided in another contribution (Léveillé-Bourret et al., in review).

**Figure 1.**
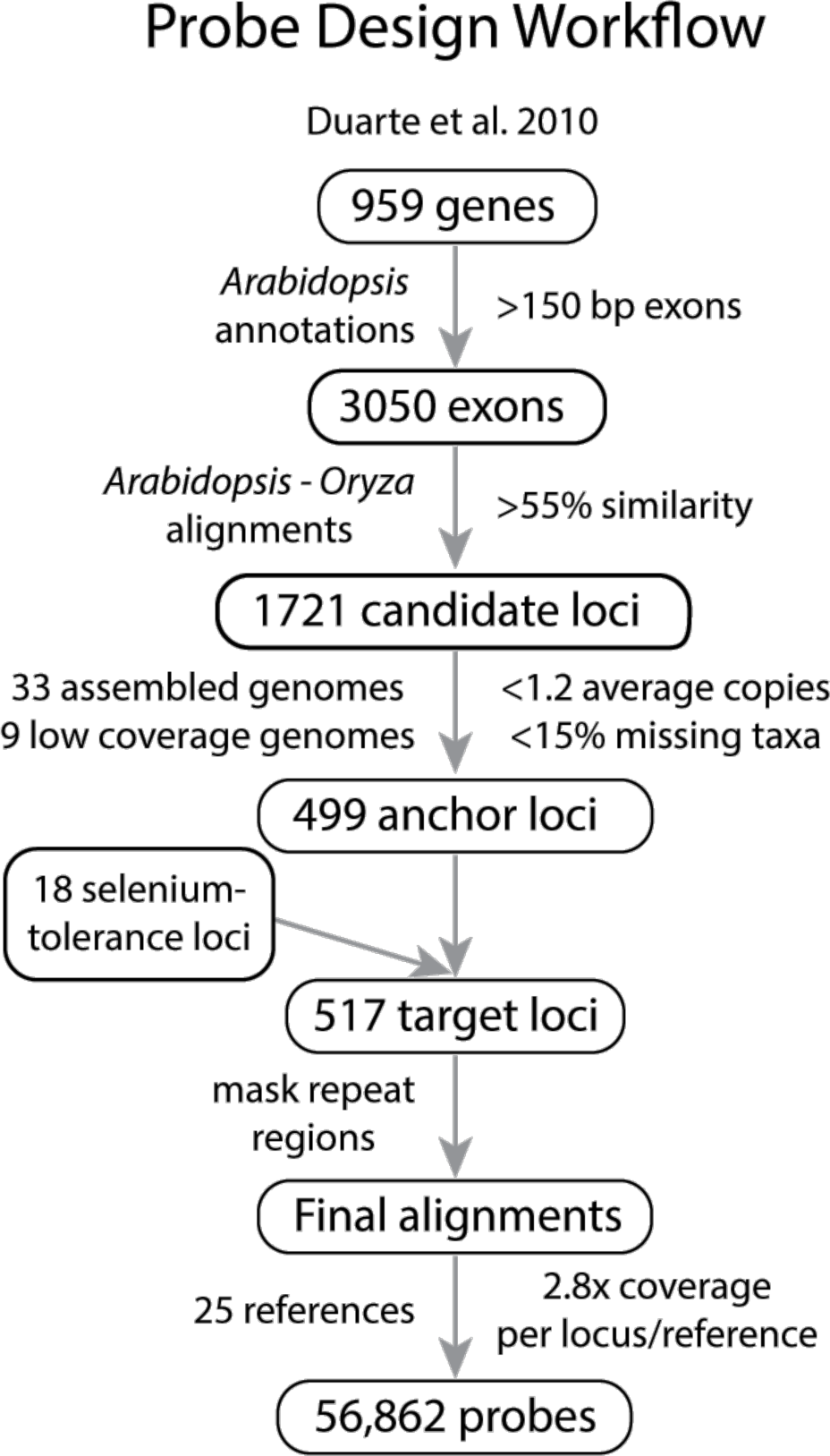
Pipeline for locus selection and probe design. Candidate loci were derived from Duarte et al. (2010) and filtered to produce anchor loci that each contained one low-copy exon with sufficient degree of conservation across angiosperms. Eighteen functional targets involved in selenium tolerance were added prior to probe tiling. Scripts used in probe design are available in Dryad (accession XXXX).

## METHODS

The primary objective of this work is to develop an enrichment kit capable of efficiently collecting a large number of informative nuclear loci across all of angiosperms. A summary of previous efforts to develop enrichment tools for plants is given in Table 1. Although a handful of previous researchers have developed enrichment kits for broad taxonomic groups (e.g., eudicots), their aim has been plastid genomes (Stull et al. 2013). Other researchers have targeted nuclear loci, but only for specific clades (e.g., *Asclepias*, Weitemier et al. 2014). Moreover, studies targeting nuclear loci have incidentally recovered plastid genomes as a byproduct of nuclear enrichment (e.g., Stephens et al. 2015b; Schmickl et al. 2016), suggesting that specifically targeting plastid genomes may be unnecessary. Our effort builds upon the effort of previous studies by developing a general angiosperm-wide enrichment resource that targets loci of moderate conservation distributed across the nuclear genome; we achieve this goal by including probes from divergent lineages representing the phylogenetic breadth of angiosperms.

### Locus Selection and Probe Design

Based on previous experience in vertebrates (Lemmon et al. 2012), including birds (Prum et al. 2015; Morales et al. 2016), snakes (Pyron et al. 2014; Ruane et al. 2015), lizards (Brandley et al. 2015; Tucker et al. 2016), fish (Eytan et al. 2015), frogs (Peloso et al. 2016) and invertebrates, including flies (Young et al. 2016), butterflies/moths (Breinholt et al. *in press*), spiders (Hamilton et al. 2016), and other taxa, development of an efficient enrichment kit that is useful across broad taxonomic scales requires careful selection of loci that are of sufficient length for probe design, relatively free of indels and introns, and of moderate conservation across the target group. In order to meet these requirements, we began with the 959 genes identified as single-copy orthologs among *Populus, Vitis, Oryza*, and *Arabidopsis* by Duarte et al. (2010). Since many of these genes contain introns, we divided these genes into exons using *Arabidopsis* annotations and retained only those that were at least 150 bp, which is long enough to contain at least two probes at 1.25 tiling density (i.e. 90 bp overlap; Fig. 1). This process yielded 3050 exons. In order to ensure we were not targeting loci that were too divergent for efficient enrichment, we removed exons with <55% sequence similarity between *Arabidopsis* and *Oryza*. The remaining 1721 exons served as our preliminary target loci.

In order to ensure that we developed enrichment probes that both represent the breadth of angiosperm diversity and also ensure that the loci captured are low in copy number across angiosperms, we leveraged a large set of published and unpublished genomic and transcriptomic resources. Specifically, we utilized complete genomes from 33 species across angiosperms (except magnoliids) together with newly obtained genomic data from nine non-model members of Poales (Table 2; Suppl. Table 1). For each of these species, we identified homologous regions following Prum et al. (2015), using *Arabidopsis* and *Oryza* as references (>55% similarity to one of the references was required). We identified the final 499 target loci as those present in >85% of the 41 references species and existing in ≤1.2 copies on average. These loci are distributed across the *Arabidopsis* genome (Suppl. Fig. 1). After selecting homologous sequences with the greatest sequence similarity to *Arabidopsis* and *Oryza* orthologs, we aligned sequences and trimmed the resulting alignments to the exon ends using the *Arabidopsis* and *Oryza* sequences as a guide. Prior to probe design, we subsampled the alignments to include only 25 species that more evenly represented angiosperm diversity. The 25 references include *Amborella*, 6 monocot families, 11 rosid families, 4 asterid families, and 4 non-rosid and non-asterid eudicot families (Table 2). Inclusion of this large number of references was necessary because enrichment success is inversely related to evolutionary distance of references and target species (Lemmon et al. 2012; Bi et al. 2013; Sass et al. 2016).

**Table 2.**
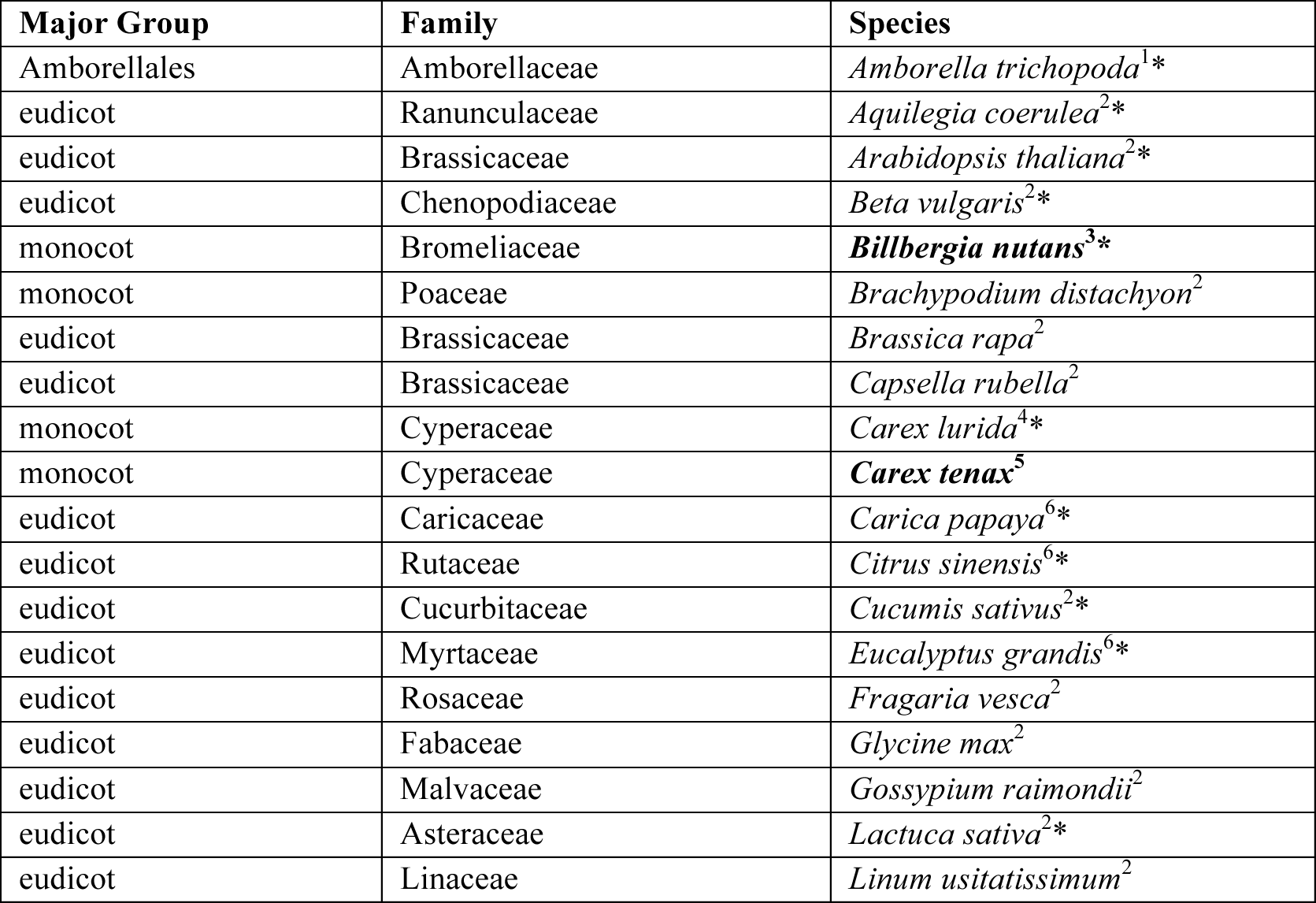
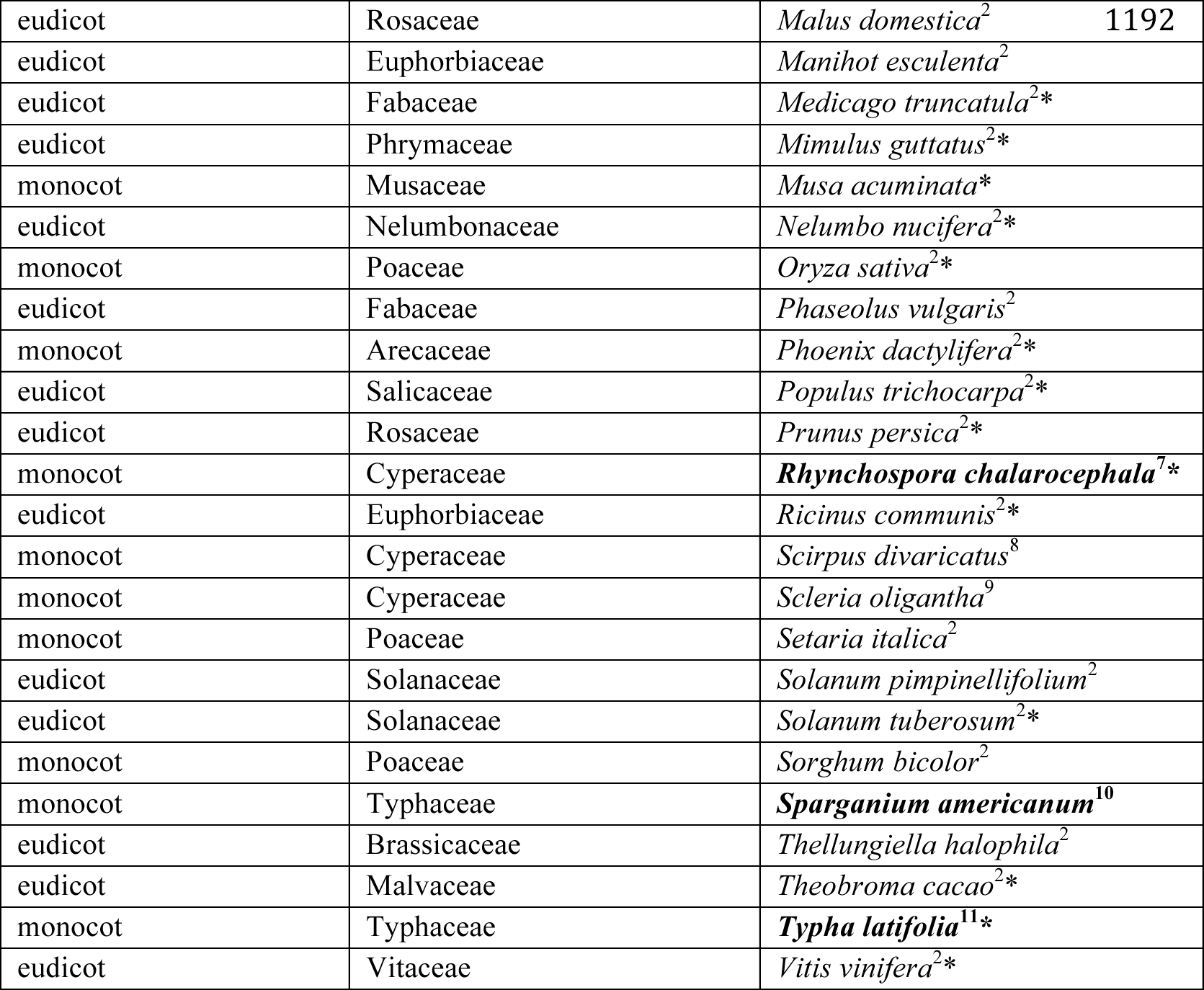
Genomic sources for taxa used in the Angiosperm v.1 probe design. All taxa included in the design (n=25) are denoted with an asterisk (*); the other taxa were utilized for locus copy number assessments only (n=18). Data for taxa indicated in bold were obtained from low-coverage genome sequencing of five vouchered specimens (Index Herbariorum code in parentheses). All other data were whole genome data downloaded from the sources provided [accessed June 20, 2013]. Sources of genomic data are indicated by superscripts as follows: 1=www.amborella.org; 2=www.phytozome.net; 3=*Chris Buddenhagen 13041601* (FSU); 4=*Loran C. Anderson 24918* (FSU); 5=*Loran C. Anderson 23871* (FSU); 6=http://bioinformatics.psb.ugent.be/plaza/; 7=*Rob Naczi 12038* (NY); 8= *Loran C. Anderson 26956* (FSU); 9= *Loran C. Anderson 26960* (FSU); 10=*Loran C. Anderson 24932* (FSU); 11=*Loran C. Anderson 23871* (FSU).

| Major Group | Family | Species |
| --- | --- | --- |
| Amborellales | Amborellaceae | *Amborella trichopoda*<sup>1*</sup> |
| eudicot | Ranunculaceae | *Aquilegia coerulea*<sup>2*</sup> |
| eudicot | Brassicaceae | *Arabidopsis thaliana*<sup>2*</sup> |
| eudicot | Chenopodiaceae | *Beta vulgaris*<sup>2*</sup> |
| monocot | Bromeliaceae | ***Billbergia nutans***<sup>3*</sup> |
| monocot | Poaceae | *Brachypodium distachyon*<sup>2</sup> |
| eudicot | Brassicaceae | *Brassica rapa*<sup>2</sup> |
| eudicot | Brassicaceae | *Capsella rubella*<sup>2</sup> |
| monocot | Cyperaceae | *Carex lurida*<sup>4*</sup> |
| monocot | Cyperaceae | ***Carex tenax***<sup>5</sup> |
| eudicot | Caricaceae | *Carica papaya*<sup>6*</sup> |
| eudicot | Rutaceae | *Citrus sinensis*<sup>6*</sup> |
| eudicot | Cucurbitaceae | *Cucumis sativus*<sup>2*</sup> |
| eudicot | Myrtaceae | *Eucalyptus grandis*<sup>6*</sup> |
| eudicot | Rosaceae | *Fragaria vesca*<sup>2</sup> |
| eudicot | Fabaceae | *Glycine max*<sup>2</sup> |
| eudicot | Malvaceae | *Gossypium raimondii*<sup>2</sup> |
| eudicot | Asteraceae | *Lactuca sativa*<sup>2*</sup> |
| eudicot | Linaceae | *Linum usitatissimum*<sup>2</sup> |

|  |  |  |  |
| --- | --- | --- | --- |
| eudicot | Rosaceae | *Malus domestica*<sup>2</sup> | 1192 |
| eudicot | Euphorbiaceae | *Manihot esculenta*<sup>2</sup> |  |
| eudicot | Fabaceae | *Medicago truncatula*<sup>2*</sup> |  |
| eudicot | Phrymaceae | *Mimulus guttatus*<sup>2*</sup> |  |
| monocot | Musaceae | *Musa acuminata*<sup>*</sup> |  |
| eudicot | Nelumbonaceae | *Nelumbo nucifera*<sup>2*</sup> |  |
| monocot | Poaceae | *Oryza sativa*<sup>2*</sup> |  |
| eudicot | Fabaceae | *Phaseolus vulgaris*<sup>2</sup> |  |
| monocot | Arecaceae | *Phoenix dactylifera*<sup>2*</sup> |  |
| eudicot | Salicaceae | *Populus trichocarpa*<sup>2*</sup> |  |
| eudicot | Rosaceae | *Prunus persica*<sup>2*</sup> |  |
| monocot | Cyperaceae | ***Rhynchospora chalarocephala***<sup>7*</sup> |  |
| eudicot | Euphorbiaceae | *Ricinus communis*<sup>2*</sup> |  |
| monocot | Cyperaceae | *Scirpus divaricatus*<sup>8</sup> |  |
| monocot | Cyperaceae | *Scleria oligantha*<sup>9</sup> |  |
| monocot | Poaceae | *Setaria italica*<sup>2</sup> |  |
| eudicot | Solanaceae | *Solanum pimpinellifolium*<sup>2</sup> |  |
| eudicot | Solanaceae | *Solanum tuberosum*<sup>2*</sup> |  |
| monocot | Poaceae | *Sorghum bicolor*<sup>2</sup> |  |
| monocot | Typhaceae | ***Sparganium americanum***<sup>10</sup> |  |
| eudicot | Brassicaceae | *Thellungiella halophila*<sup>2</sup> |  |
| eudicot | Malvaceae | *Theobroma cacao*<sup>2*</sup> |  |
| monocot | Typhaceae | ***Typha latifolia***<sup>11*</sup> |  |
| eudicot | Vitaceae | *Vitis vinifera*<sup>2*</sup> |  |

In order to test the potential for simultaneously targeting loci of known function in addition to anchor loci, we incorporated probes targeting 18 selenium-tolerance genes. The 18 genes are candidates for the selenium-tolerance phenotype obtained from a comparative transcriptome analysis between selenium hyperaccumulator *S. pinnata* and selenium non-hyperaccumulator *S. elata* (unpublished data) and a macroarray study between *S. pinnata* and *S. albescens* (Freeman et al. 2010). Selenium is atomically similar to sulfur and can be assimilated into selenocysteine via the sulfur assimilation pathway (Terry et al. 2000). These genes include sulfur transporters and key enzymes in sulfur assimilation. Alignments for 18 of these genes were obtained from TAIR (Berardini et al. 2015). Following the methods described above, we scanned genomic resources for the 25 references species for selenium homologs using *Arabidopsis* as a reference. Alignments of the homologs were then constructed using MAFFT (v7.023b; Katoh and Standley 2013) and trimmed as described above.

After masking high-copy and repetitive regions, we tiled 120 bp probes across each of the sequences at 2.8x coverage (neighboring probes overlap by approximately 48 bp). Probes were synthesized by Agilent Technologies (Santa Clara, CA) to develop the kit hereafter referred to as the Angiosperm v.1 design. The alignments and probe design are available as Supplemental Material (Dryad accession XXXX).

### Taxon Sampling

Data were collected from a total of 104 angiosperm species representing 30 orders and 48 families (Suppl. Table 2). For 53 of these species (representing six orders, 10 families, and 21 genera), the Angiosperm v.1 kit was used to enrich genomic DNA for the target regions prior to sequencing. Sequences for the remaining 51 species were obtained from low-coverage genomic reads (not enriched), assembled transcriptomes, and assembled genomes (Suppl. Table 2).

### Hybrid Enrichment Data Collection

In order to test the efficiency and utility of the AHE resource described above, we enriched 53 samples from across the phylogenetic breadth of angiosperms following the general methods of Lemmon et al. (2012), with adaptations made for plant samples. Genomic DNA was extracted using the DNEasy Plant Mini Kit (Qiagen, Valencia, California, USA). The protocol was modified following suggestions (Costa and Roberts 2014) to compensate for frequent low yields obtained with the standard protocol. After extraction, genomic DNA was sonicated to a fragment size of ~300-800 bp using a Covaris E220 Focused-ultrasonicator with Covaris microTUBES. Subsequently, library preparation and indexing were performed on a Beckman-Coulter Biomek FXp liquid-handling robot following a protocol modified from Meyer and Kircher (2010). A size-selection step was also applied after blunt-end repair using SPRI select beads (Beckman-Coulter Inc.; 0.9x ratio of bead to sample volume). Indexed samples were then pooled at approximately equal quantities (typically 16-18 samples per pool), and then each pool was enriched using the Angiosperm v.1 kit (Agilent Technologies Custom SureSelect XT kit ELID 623181). After enrichment, 3–4 enrichment reactions were pooled in equal quantities for each sequencing lane and sequenced on PE150 Illumina HiSeq 2500 lanes at the Translational Science Laboratory in the College of Medicine at Florida State University.

### Paired-Read Merging

In order to increase read accuracy and length, paired reads were merged prior to assembly following Rokyta et al. (2012). In short, for each degree of overlap each read pair was evaluated with respect to the probability of obtaining the observed number of matches by chance. The overlap with the lowest probability was chosen if the p-value was less than 10^−10^. This low p-value avoids chance matches in repetitive regions. Read pairs with a p-value below the threshold were merged and quality scores were recomputed for overlapping bases (see Rokyta et al. [2012] for details). Read pairs failing to merge were utilized but left unmerged during the assembly.

### Read Assembly

Divergent reference assembly was used to map reads to the probe regions and extend the assembly into the flanking regions (Fig. 2; Prum et al. 2015). More specifically, a subset of the taxa used during probe design were chosen as references for the assembly: *Arabidopsis thaliana, Billbergia nutans*, and *Carex lurida*. Matches were called if 17 bases matched a library of spaced 20-mers derived from the conserved reference regions (i.e., those used for probe design). Preliminary reads were then considered mapped if 55 matches were found over 100 consecutive bases in the reference sequences (all possible gap-free alignments between the read and the reference were considered). The approximate alignment position of mapped reads were estimated using the position of the spaced 20-mer, and all 60-mers existing in the read were stored in a hash table used by the de-novo assembler. Simultaneously using the two levels of assembly described above, the three read files were traversed repeatedly until a pass through the reads produced no additional mapped reads. For each locus, a list of all 60-mers found in the mapped reads was compiled and 60-mers were clustered if found together in at least two reads. Contigs were estimated from 60-mer clusters. In the absence of contamination, low coverage, and gene duplication each locus should produce one assembly cluster. Consensus bases were called from assembly clusters as unambiguous base calls if polymorphisms could be explained as sequencing error (assuming a binomial probability model with the probability of error equal to 0.1 and alpha equal to 0.05). Otherwise ambiguous bases were called (e.g., ‘R’ was used if ‘A’ and ‘G’ were observed). Called bases were soft-masked (made lowercase) for sites with coverage lower than five. Assembly contigs derived from less than 10 reads were removed in order to reduce the effects of cross contamination and rare sequencing errors in index reads. Scripts used in the assembly process are available on Dryad (accession XXXX).

**Figure 2.**
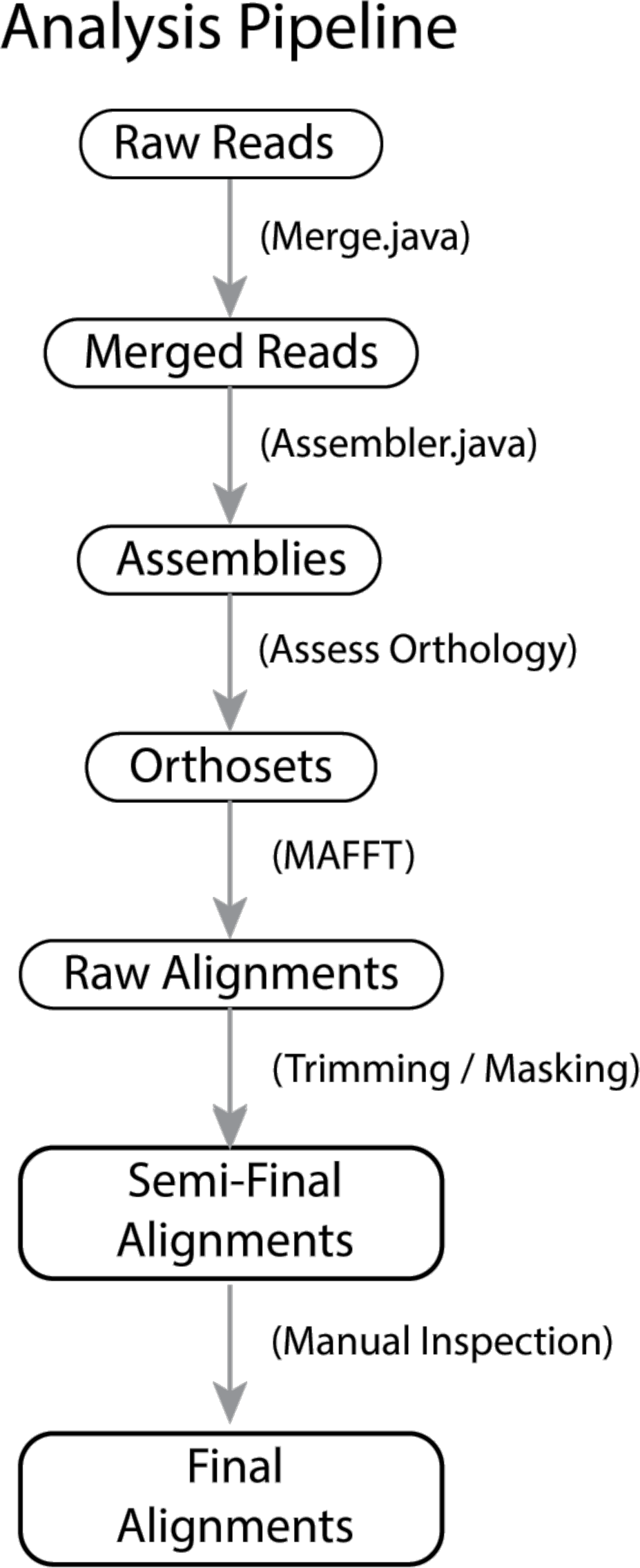
Standard pipeline for processing anchored phylogenomic data to produce final alignments. Merged read pairs are assembled using a quasi-de novo approach that utilizes divergent references from the kit. After orthologous sets of consensus sequences are identified, raw alignments are constructed using MAFFT and trimmed/masked to remove probable misaligned regions. Alignments are finalized after being manually inspected and trimmed in Geneious. Scripts are available in Dryad (accession XXXX).

### Orthology Estimation

After grouping homologous sequences obtained by enrichment and whole genomes, putative orthologs were identified for each locus following Prum et al. (2015). For each locus, pairwise distances among homologs were computed using an alignment-free approach based on percent overlap of continuous and spaced 20-mers. Using the distance matrix, sequences were clustered using the neighbor-joining algorithm, but allowing at most one sequence per species to be present in a given cluster. Note that flanks recovered through extension assembly contain more variable regions and allow gene copies to be sorted efficiently. Gene duplication before the ancestor of the clade results in two distinct clusters that are easily separated. Duplication within the clade typically results in two clusters, one containing all of the taxa and a second containing a subset of the taxa (missing data). Gene loss also results in missing data. In order to reduce the effects of missing data, clusters containing fewer than 50% of the species in the taxon set were removed from downstream processing.

### Alignment and Trimming

Putatively orthologous sequences were processed using a combination of automated and manual steps in order to generate high quality alignments in a reasonable amount of time. Sequences in each orthologous cluster were aligned using MAFFT v7.023b (Katoh and Standley 2013), with --genafpair and --maxiterate 1000 flags utilized. The alignment for each locus was then trimmed/masked using the steps from Prum et al. (2015). First, each alignment site was identified as “conserved” if the most commonly observed character state was present in > 40% of the sequences. Second, using a 20 bp window, each sequence was scanned for regions that did not contain at least 10 characters matching to the common state at the corresponding conserved site. Characters from regions not meeting this requirement were masked. Third, sites with fewer than 12 unmasked bases were removed from the alignment. A visual inspection of each masked alignment was carried out in Geneious version 7 (www.geneious.com; Kearse et al. 2012). Regions of sequences identified as obviously misaligned or non-homologous were removed.

### Phylogeny Estimation

Adequate modeling of sequence evolution is an important prerequisite for accurate phylogenetic estimation (Hillis et al. 1994; Yang 1994; Posada and Crandall 1998; Lemmon and Moriarty 2004; Ripplinger and Sullivan 2008; Lanfear et al. 2012; Hoff et al. 2016). Site partitioning is one aspect of modeling that has recently gained appreciation (Brown and Lemmon 2007; Lanfear et al. 2012; Darriba and Posada 2015; Frandsen et al. 2015). When the loci being modeled are from protein coding regions, some authors choose to partition by codon position and remove third codon positions in order to reduce the effects of substitutional saturation (e.g., Breinholt and Kawahara 2013; Wickett et al. 2014; Frandsen et al. 2015). Although the anchored loci we target here were derived from protein-coding exonic regions, partitioning by coding position would not be sufficient because captured flanks could contain intronic and intergenic regions that are also likely to vary both by model and rate of evolution. Moreover, the degree of saturation and its effects on downstream analyses depend heavily on the phylogenetic breadth and depth of sampling. Consequently, we used an agnostic approach to site partitioning that does not rely on a prior knowledge of possible site partitions. More specifically, we used PartitionFinder v2.0.0-pre14 (Frandsen et al. 2015), which utilizes a k-means approach to partition sites into clusters based on similarity with respect to model of sequence evolution (including rate). This approach produces partitioning schemes that fit alignments better than schemes determined by methods requiring definition of *a priori* partitions (e.g., by codon positions and locus boundaries; Frandsen et al. 2015). An added benefit to using this approach is that it produces estimates of site-specific rates of evolution.

In order to allow for estimation of trees under both coalescent and supermatrix approaches, RAxML v8.1.21 (Stamatakis 2014) was used to estimate gene trees from individual locus alignments and also the concatenated supermatrix (GTRGAMMA model, partitioned as described above, with -f -a flags). In each case, *Amborella* was specified as the outgroup, and 100 rapid bootstrap replicates were collected in order to measure support for the maximum likelihood estimate. All other parameters remained at default settings. Species trees were estimated in ASTRAL-II v.4.9.7 (Mirarab and Warnow 2015) using bootstrap replicates from the RAxML-estimated gene trees.

### Evaluating the Size and Variability of Flanking Regions for Family-and Genus-Level Clades

One feature of AHE is that data collected by the approach can be useful at both deep, intermediate, and shallow timescales. Recall that in order to ensure efficient enrichment across deep time scales, regions targeted by the enrichment kit described above consist of moderately conserved exons. Consequently, flanking regions are expected to contain intronic and intergenic sites that may only be accurately aligned at shallower taxonomic scales. We expect, therefore, that the majority of flanking bases would be filtered out when the angiosperm-wide alignment is trimmed. In order to study the effect of phylogenetic depth on the size of usable flanks, we also performed the post-assembly analysis steps on subsamples of the taxa. More specifically, we performed orthology assessment and constructed trimmed alignments specific to the following clades: Cyperaceae (21 samples), Caryophyllales (5 samples), Dipsacales (6 samples), *Magnolia* (7 samples), *Protea* (4 samples), and *Stanleya* (5 samples). The size and variation of flanking regions for these alignments was compared to corresponding sections of the angiosperm-wide alignment.

### Assessing Robustness of Phylogenetic Estimates

Phylogenetic estimates can be highly sensitive to not only the model of evolution assumed, but also the choice of data being analyzed (Ripplinger and Sullivan 2008; Wickett et al. 2014; Hosner et al. 2015; Simmons et al. 2016). Inclusion of saturated sites and/or outlier loci, for example, can have particularly profound effects on phylogenetic estimates (Breinholt and Kawahara 2013; Simmons et al. 2016). These effects can result in conflict among studies, especially for phylogenetic relationships that are particularly difficult (e.g., the relationships among eudicots, monocots, and magnoliids). Given this difficulty, it is surprising that more authors do not present results from a more exhaustive exploration of the effects of their data selection on their estimates of phylogeny.

Here, we present an approach to evaluate the robustness of phylogenetic estimates to differential character and taxon sampling (Fig. 3; all scripts available in Dryad). Although we focus on the effects of variation in sampled loci included, other aspects of the data set and analysis could also be incorporated (e.g., taxon sampling). The approach begins by independently varying two or more aspects of the data set or analysis. In this case we varied both the sites included at each locus and also the loci included. We ordered the sites based on evolutionary rate and the loci based on similarity to other loci estimated via gene tree distances; then we analyzed the data with successively less variable and less distant loci included (see details below). Data sets generated under each of these conditions were then used to estimate the phylogeny and support values inferred using both supermatrix and coalescent approaches. The novel aspect of this approach is the representation of the resulting support values for particular nodes as heat maps in the parameter space defined by the aspects of the data set or analysis that we varied.

**Figure 3.**
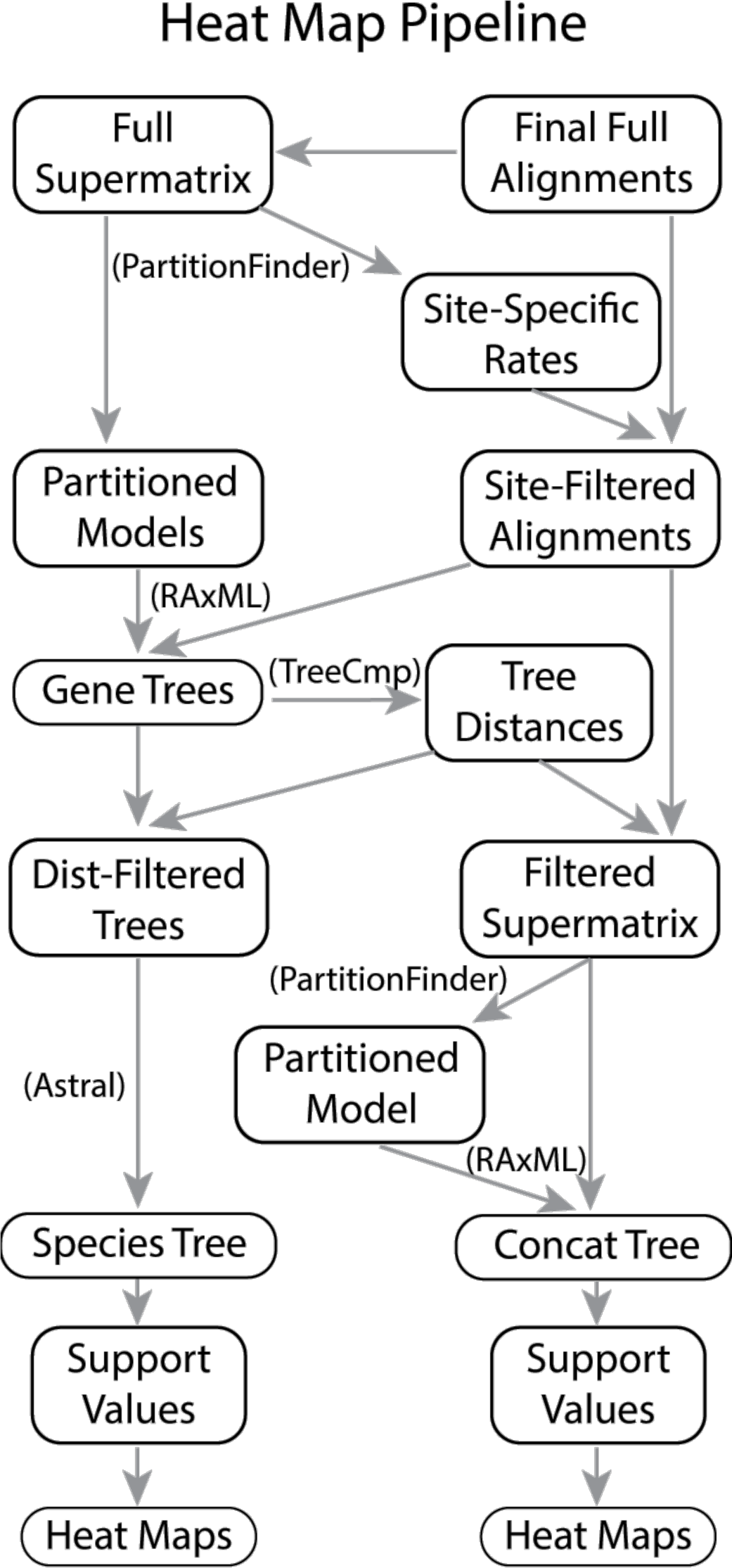
Pipeline for generating heat maps representing robustness of phylogenetic estimates to variation in site and locus filtering. Labeled arrows involve steps performed using standard methods, whereas unlabeled arrows involve steps performed using custom scripts available in Dryad (accession XXXX).

High evolutionary rate estimates can result from alignment error, substitutional saturation, and convergent evolution of base composition. Removal of the sites with highest estimated rates can improve the support for the topology by removing noise from the data set (Castresana 2000; Talavera and Castresana 2007; Xia and Lemey 2009). Removing too many sites, however, will eventually result in loss of signal and subsequently in poorly supported topologies. In some cases, conversely, the most rapidly evolving sites contain strong phylogenetic signal, and so removing these sites may actually reduce branch support values. Given the difficulty in knowing *a priori* which rate threshold optimizes this tradeoff for a particular data set, a useful approach is to test several thresholds along a continuum. We utilized site-specific rates estimated by PartitionFinder v2.0.0-pre14 from a concatenated matrix containing all loci using the kmeans option (default settings otherwise; Frandsen et al. 2015). Fourteen site inclusion thresholds along the observed distribution were then chosen based on the following percentiles: 100%, 99%, 97%, 95%, 92%, 89%, 86%, 82%, 76%, 70%, 63%, 54%, 43%, and 31%. These thresholds progressively remove sites such that our test would be more sensitive to high levels of site inclusion. Based on preliminary tests with this data set, we expected phylogenetic support to deteriorate rapidly when less than 63% of sites were included. New locus-specific alignments were then constructed based on each of these site inclusion levels, and gene trees were estimated for each of these data sets using RAxML as described above (14 estimated gene trees per locus; Fig. 4a).

**Figure 4.**
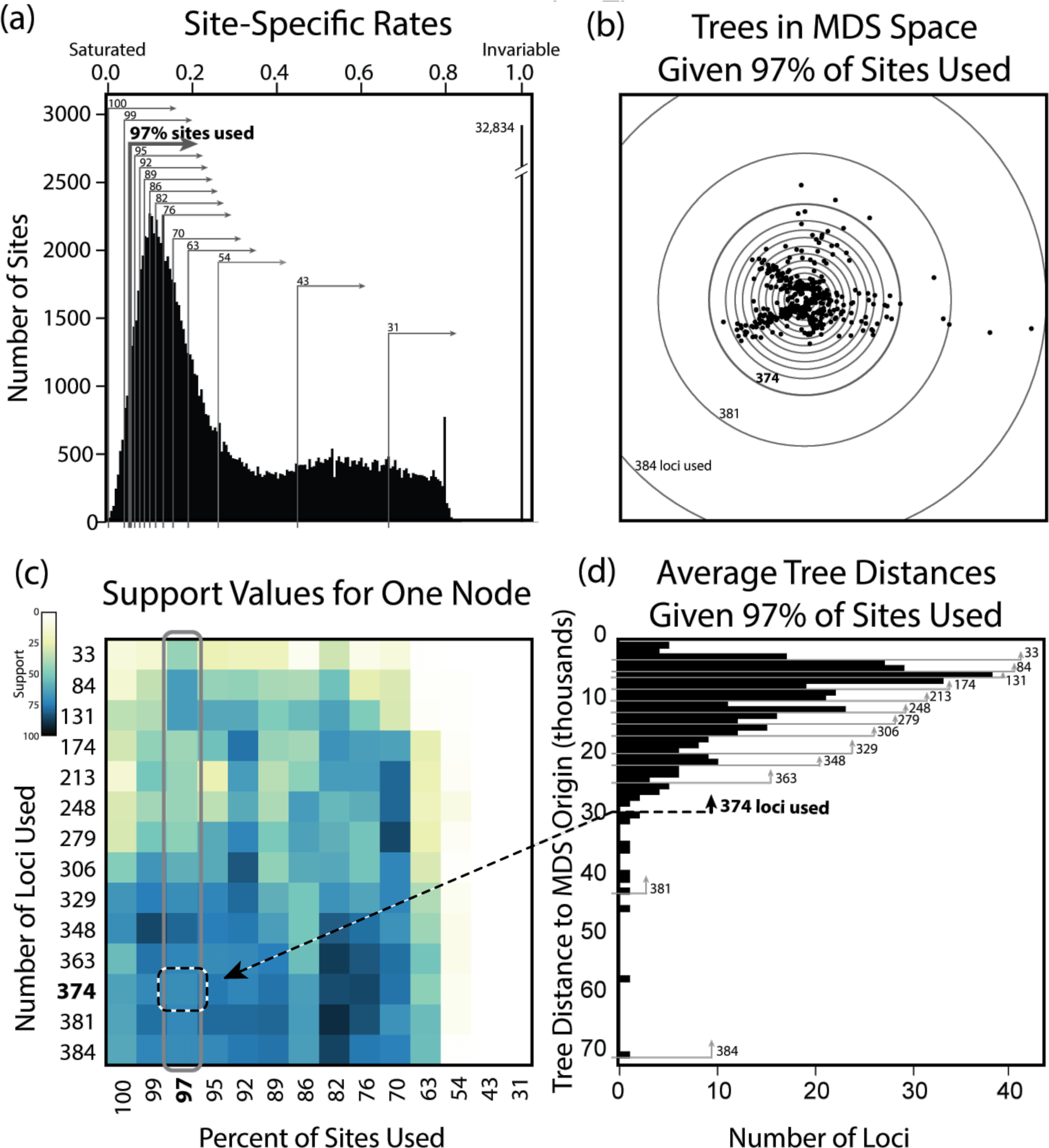
Strategy for generating 196 data filtering conditions to assess robustness of phylogenetic estimates. (a) Sites are ranked by rate as estimated using kmeans in PartitionFinder. Invariable sites are plotted to right of the distribution. Fourteen site-inclusion thresholds (gray lines/arrows on histogram) are applied, with the most variable sites being excluded at the lowest thresholds. Gene trees constructed after applying each of these thresholds are compared pairwise using the triplet measure in TreeComp to produce a pairwise tree-distance matrix. (b) Plotting loci in multidimensional space (MDS) using corresponding tree-distances allows for the identification of outliers (furthest from center) and 14 thresholds for inclusion of loci for phylogeny estimation (gray circles). (c) Example of heat-map for one node showing phylogenetic support as color for each of the 14×14=196 combinations/data sets, with white indicating no support for the node and black indicating 100% support for the node. (d) Thresholds (gray lines/arrows) plotted on the distribution of loci ranked by distance from origin on the MDS plot. Robust estimates show relatively consistent level of support (similar color) across the heat map in (c).

Inclusion of loci with insufficient signal can result in imprecise or inaccurate estimates of species trees (Hosner et al. 2015; Simmons et al. 2016). Moreover, inclusion of loci with estimated gene-tree-histories that are substantially different than the underlying species history can also result in imprecise or inaccurate estimates of species trees (Meredith et al. 2011; Townsend et al. 2011; Simmons and Gatesy 2015; Simmons et al. 2016; Springer and Gatesy 2016). Therefore, removal of outlier gene trees may result in more accurate and precise species trees. As with inclusion of sites, however, inclusion of too many loci will eventually result in reduced accuracy, precision, and support. In order to identify the level of locus inclusion that maximizes accuracy and precision, we tested several thresholds along a continuum. For each of the 14 sets of gene trees described above, pairwise tree distances were estimated using treeCmp (v1.0-b29; Bogdanowicz et al. 2012), via the triple metric for rooted trees (tt option; Critchlow et al. 1996). The resulting matrix of pairwise tree distances was visualized using multidimensional scaling using the R package plotrix version 3.6-2 (Fig. 4b; (Fig. 3; from Lemon [2006]). Euclidian distance from each tree to the center of the trees was then used to rank the loci, with the greatest distance (i.e., greatest outlier) having the highest rank. Fourteen locus-inclusion thresholds were then applied: 384, 381, 374, 363, 348, 329, 306, 279, 248, 213, 174, 131, 84, and 33 loci. These thresholds were chosen to cover a broad range of locus inclusion levels, but still allow the effects of removing small numbers of outliers to be studied. These 14 locus-inclusion thresholds were applied to each of 14 site-inclusion thresholds described above (Fig. 4d). Phylogenies were then inferred using each of the 196 combinations in RAxML and ASTRAL-II, using corresponding concatenated matrices and gene trees sets, respectively.

The primary goal of the extensive analyses described above is to produce a mechanism for visualizing the sensitivity of phylogenetic estimates to variation in site and locus inclusion. To this aim, nodal support for trees at each filter level were estimated using the R package ape (Paradis et al. 2004) with the “prop.clades” command, which counts the number of times the bipartitions present in a reference tree are present in a list of trees. After selecting a reference tree (the best tree from the full unfiltered data set), the occurrence of a clade in the bootstrap trees was assessed for each of the 196 ASTRAL-II and 196 RAxML concatenated analyses. This step produced support values for each combination of thresholds at each node in the reference tree. Heat maps were then constructed for each internal node in the reference tree (Fig. 4c). Note that alternate reference trees can be used to generate heat maps representing support for alternative hypotheses.

### Testing Alternative Hypotheses

The large number of estimates produced by the above procedure provides an opportunity to test alternative phylogenetic hypotheses and alternative phylogeny estimation approaches across a broad range of conditions. To demonstrate the potential for testing alternative phylogenetic hypotheses, we focused on the relationship between eudicots, monocots, and magnoliids that has been debated in the literature (Lee et al. 2011; Soltis et al. 2011; Burleigh et al. 2011; Wickett et al. 2014). Specifically, support values represented by heat maps corresponding to support for the three alternative relationships among these clades (magnoliids-monocots sister, eudicots-monocots sister, and magnoliids-eudicots sister) were compared using randomization tests as follows. Average support across all analysis conditions (a heat map) was computed for each of the three relationships. A test statistic was then computed as the difference between the mean support (across analysis conditions) for two alternative relationships. To generate a null distribution with which to compare the test statistic, support values found in heat map matrices were then randomized the relevant pair of matrices, while keeping the analysis condition constant. In other words, the two support values (one for each relationship) for a given analysis condition were randomized with respect to the matrix in which they were placed. After randomizing, the difference between the means of the two randomized matrices was computed as a value in the null distribution. This process was repeated 10,000 times to generate the null distribution. To demonstrate the potential for testing support given by alternative analysis methods, we used the randomization test described above, but instead used matrices derived from the supermatrix (RAxML concatenated) and species tree (ASTRAL-II) analyses.

We also tested the hypothesis that support produced by the species tree approach is more robust to variation in data sets than the support produced under the supermatrix approach (Song et al. 2012; Edwards et al. 2016). In particular, we compared the overall sensitivity as the variance in support values across analysis conditions, averaging the variance across nodes (a value of zero indicates high robustness). We then computed the test statistic as the difference between the sensitivity measure for the coalescent and supermatrix (with supermatrix - coalescent > 0 indicating support that the coalescent approach is more sensitive to changes in analysis conditions). Ten thousand points from the null distribution were computed by shuffling corresponding support values between coalescent-and supertree-derived matrices, before recomputing the test statistic.

In addition to the randomization tests described above, we performed more traditional hypothesis tests using point estimates of phylogeny from the analyses of the full data set. The best-constrained trees for each topology were estimated in RAxML using the -g option under a GTRGAMMA model. The per-site log likelihoods for the three best constrained trees were written to file and loaded in the CONSEL (v0.1i) program (Shimodaira and Hasegawa 2001). This program ranks alternative hypotheses in order of the likelihood via p-value calculated from the multi-scale bootstrap using several tests including the widely used Shimodaira–Hasegawa (SH) and the approximately unbiased (AU) test (Shimodaira and Hasegawa 2001; Shimodaira 2002).

## RESULTS

### Locs Selection and Probe Design

The procedure followed for locus selection (Fig. 1) yielded a large number of target loci, with minimal missing data and optimal levels of sequence variation for hybrid-enrichment-based phylogenomics. Target probes were designed for 499 anchor loci and 18 functional loci (517 loci in total). Alignments of sequences for the 25 reference species averaged 343 sites per target locus (range: 124 to 2176 sites) and contained only 10% missing characters. Sequence variation in target regions was moderate: 62% of the sites were variable, 51% of the sites were parsimony informative, pairwise sequence divergence averaged 27%, and maximum pairwise sequence divergence within loci averaged 39%. The probe design contained a total of 56,862 probes (~4.4 probes per species per locus; See Supplemental Materials).

### Hybrid Enrichment

The probe kit developed above performed well on the 53 test samples taken from across angiosperms; >90% of target loci were obtained for 90% of the samples (Table 3). Over 91 billion nucleotides were sequenced on the two PE150 Illumina lanes. Assemblies resulted in recovery of an average of 470 loci (91%) with contigs longer than 250 bp. The average consensus sequence recovered was 764 bp, more than twice the length of average probe region (343 bp). On average, 14% of reads mapped to target plus flanking regions (i.e., were incorporated in the assemblies). As expected, capture efficiency was lower for taxa more evolutionary divergent from reference sequences (Fig. 5). For example, an average of only 329 loci were recovered for members of Piperales, which shares an ancestor with the closest reference 124 million years ago (Magallón et al. 2013). In contrast, for members of Brassicales (sharing a common ancestor with the closest reference 24 million years ago), an average of 512 loci were recovered.

**Figure 5.**
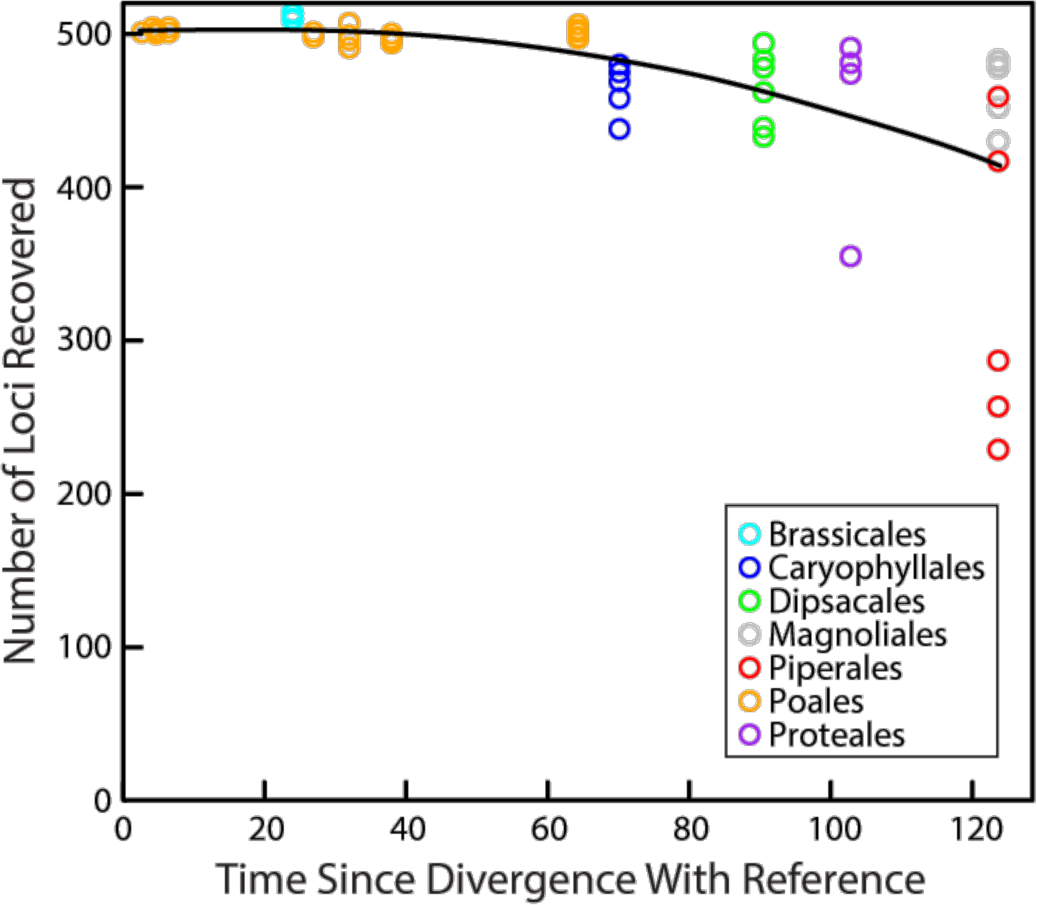
The relationship between locus recovery and taxonomic representation in the enrichment kit. For each enriched sample, the number of loci recovered with consensus sequence greater than 250 bp is plotted against the divergence time (in millions of years - see Supplemental Methods for details) between that sample and the nearest reference species used for probe design. The curve represents the best-fit quadratic polynomial (y = -0.0083x^2^ + 0.3049x + 501.43; R^2^ = 0.35734).

**Table 3.**
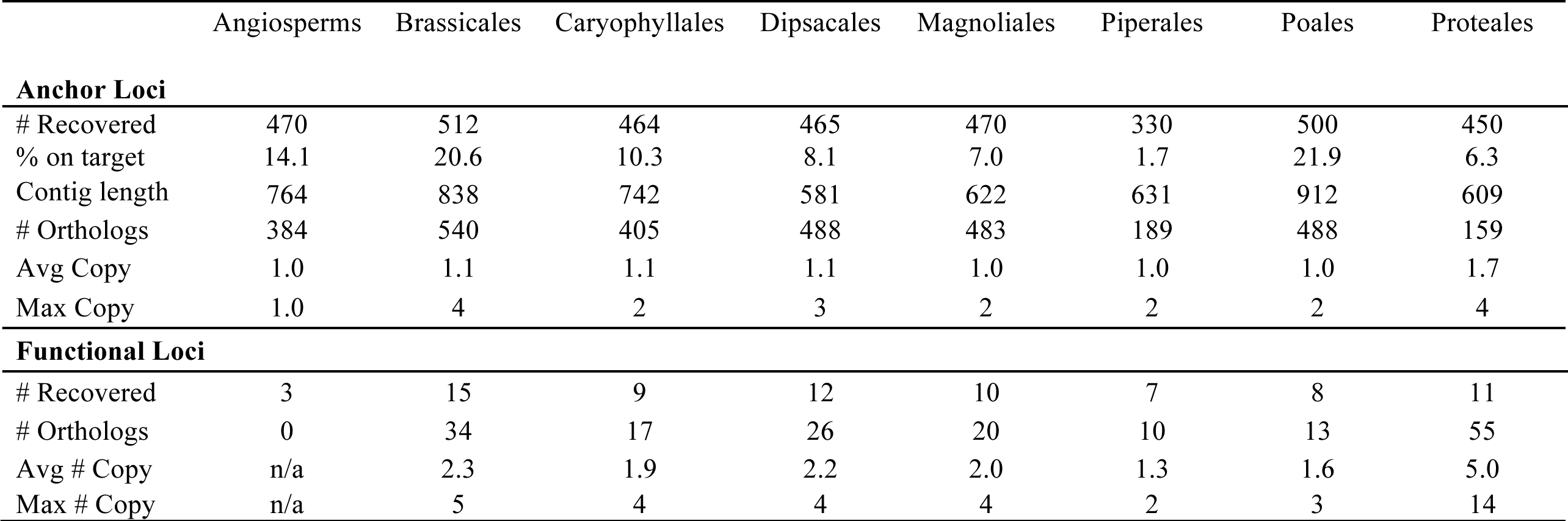
Overall success for targeted loci, in terms of regions remaining in the alignment for phylogenetic estimation after orthology and trimming phases of the bioinformatics pipeline were completed. Targets include the 499 low-copy nuclear loci and the 18 functional mostly high-copy genes associated with selenium tolerance. The total number of orthologs indicates the total number of loci in the alignment, which may be larger than the number of targets since some targets appear to be multicopy.

### Alignments

For the angiosperm taxon set, a large number of loci were retained through the post-assembly pipeline. More specifically, 384 alignments containing orthologous sequences were obtained (available as Supplemental Material). The final concatenated angiosperm alignment contained 138,616 bases. In general, alignments contained a modest amount of missing data (7% if missing taxa were excluded, 22% if missing taxa were included). The number of missing taxa varied among loci (mean=15, range=2-40). Note for the angiosperm-wide alignments, flanks were typically trimmed off because they contained non-homologous or unalignable bases at this deep scale (see below for intermediate-and shallow-clade alignments). As these alignments contain primarily probe regions, it is not surprising that final, trimmed alignments were only slightly longer than the alignments used to design probes (360 vs. 343, respectively). Also note that none of the 18 functional loci were retained for the angiosperm-wide taxon set.

For the intermediate and shallow-clade taxon sets, an even larger number of loci was retained through the post-assembly pipeline (Table 3). An average of 410 alignments containing orthologous sequences were obtained (range 189 to 540). Because they contained flanking regions, alignments for these genus-level taxon sets were substantially longer (68% longer on average) than those of the angiosperm-level taxon set (Table 4). Concatenated alignments averaged 233,993 bases (107,918 to 365,941), and locus-specific alignments averaged 577 bases (375 to 670). Remarkably, all taxa were present for each of the final locus sets, indicating that missing data/taxa in the angiosperm-wide alignments resulted from differences in locus recovery across families. One indication of this cause is the consistent absence of some target loci in Piperales and Proteales, in which only 189 and 159 orthologous loci were retained, respectively. As a result of the increased length and variability of the loci, support for the intermediate and shallow-scale clades improved (Suppl. Figs. 2-6). Although the estimated topologies remained unchanged, half of the nodes that did not receive full support (100% bootstrap) in the Angiosperm-wide alignment became fully supported by the intermediate and shallow-scale alignments.

**Table 4.** Properties of probe regions and flanks for deep, intermediate, and shallow-scale alignments.

|  | Probe Region |  |  |  | Flanks |  |  |  |
| --- | --- | --- | --- | --- | --- | --- | --- | --- |
|  | Total Sites | Var. Sites | Inform. Sites | % Var. | Total Sites | Var. Sites | Inform. Sites | % Var. |
| Angiosperms | 138,616 | 103,616 | 94,185 | 74.8 | N/A | N/A | N/A | N/A |
| *Carex* | 114,623 | 16,512 | 3,592 | 13.9 | 188,315 | 43,195 | 11,307 | 23.6 |
| Caryophyllales | 96,372 | 23,355 | 9,171 | 23.6 | 66,471 | 19,490 | 6,964 | 29.2 |
| Cyperaceae | 120,774 | 25,082 | 7,990 | 19.8 | 224,529 | 70,104 | 20,716 | 31.6 |
| Dipsacales | 80,064 | 21,501 | 10,151 | 26.3 | 12,429 | 3,166 | 1,315 | 23.4 |
| *Magnolia* | 110,509 | 6,460 | 1,181 | 5.6 | 87,905 | 10,450 | 1,843 | 12.4 |
| *Protea* | 82,258 | 11,391 | 544 | 13.7 | 66,269 | 4,644 | 536 | 7.3 |
| *Rhynchospora* | 132,656 | 29,530 | 16,615 | 20.9 | 164,027 | 60,759 | 35,824 | 36.9 |
| *Stanleya* | 129,500 | 2,036 | 149 | 1.4 | 178,670 | 11,933 | 1,975 | 6.9 |

One reason why a greater number of orthologous loci were retained for the intermediate and shallow taxon sets is that orthology could more easily be established for duplicated genes. For example, most of the functional locus targets contained multiple copies (average 2.16; Table 3). At one functional target locus in Proteaceae, in fact, 14 orthologs were identified. Multi-copy loci such as these were removed from the angiosperm-wide taxon set during the orthology/trimming process because copy number for these loci varied wildly across genera (i.e., multiple independent duplications resulting in these copies occurred within angiosperms). Brassicales, in which many species have high selenium tolerance (likely due to the propensity for sulfur accumulation and assimilation into unique sulfur-containing compounds), produced the greatest number of orthologous alignments (n=574).

### Robustness of Phylogenetic Estimates

Although the majority of the angiosperm relationships were unequivocally supported across the majority of analysis conditions, some relationships were less robust to variation in levels of data filtering and/or method of phylogeny estimation. The robustness of each estimated node is shown as a heat map in Figure 6, which provides not only a comprehensive view of support across many angiosperms orders, but also a clear indication of which relationships are only supported under specific conditions. Many nodes were well supported in all but the most extreme thresholds for site and locus removal (i.e., >60% of sites removed and/or >60% loci removed). Overall, support values were consistent between coalescent and supermatrix approaches as long as more than ~100 loci were included. Nonetheless, support for several nodes varied wildly across analysis conditions, with highest support occurring when a moderate to large numbers of sites or loci were removed.

**Figure 6.**
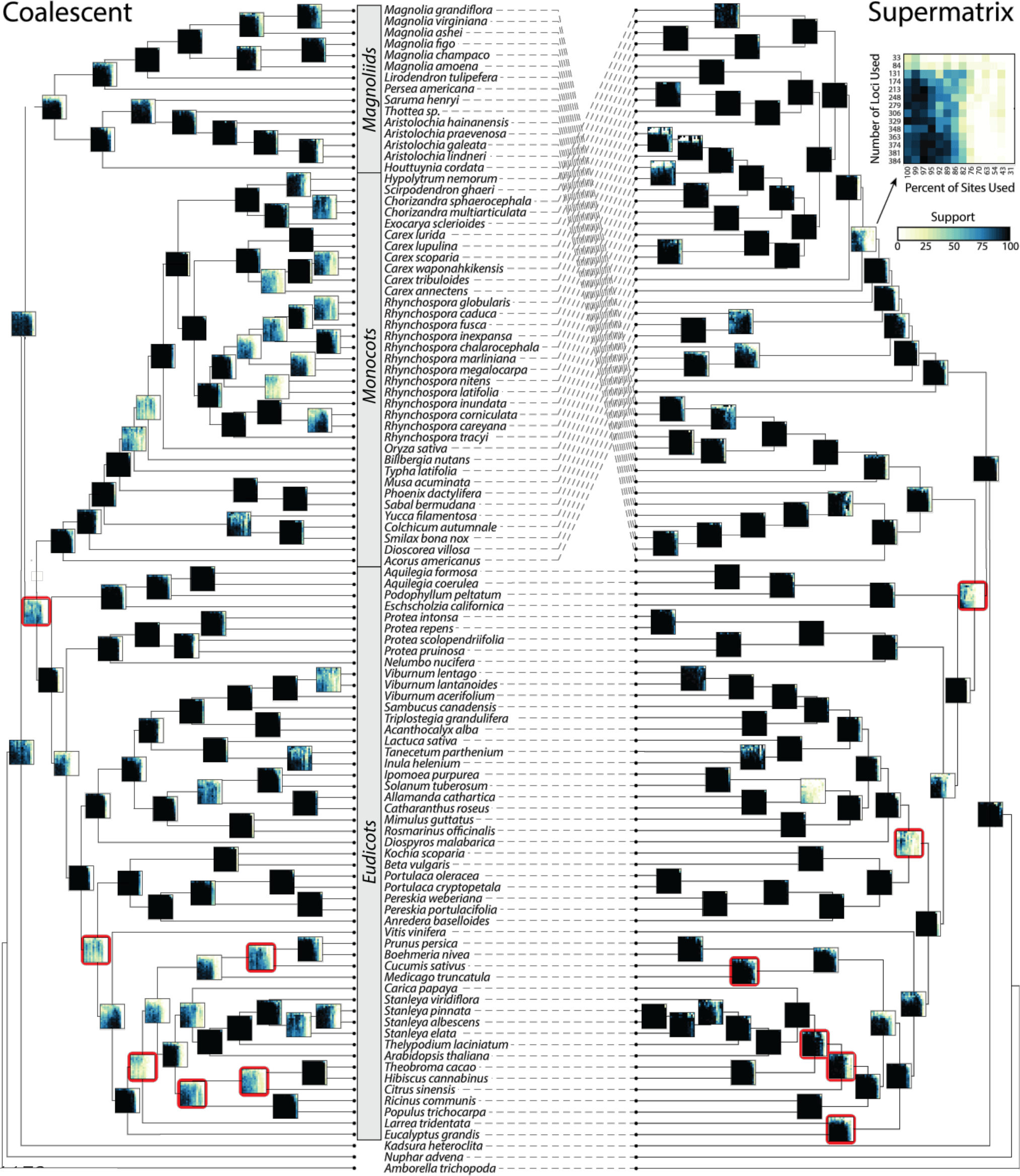
Estimates of the angiosperm phylogeny under a coalescent (ASTRAL) or supermatrix (concatenated RAxML) framework with all data included. Heat-maps on each internal node indicate bootstrap support across 196 data inclusion conditions (example shown in upper right), with lower-left corner of each heat map indicating support when all data are included (colors described in Fig. 4). Solid black heat maps indicate high support for a node under a broad range of conditions. Heat maps for nodes not present in both topologies are outlined in red.

For the relationship of particular interest, that between monocots, eudicots and magnoliids, support for alternative arrangements were highly dependent upon the level of site filtering, the level of locus filtering, and also the approach to phylogeny estimation (Fig. 7). Coalescent-based analyses supported the eudicots-monocots sister hypothesis with moderate to high support for the majority of site-and locus-filtering conditions. Support for the other two arrangements was almost entirely lacking, regardless of the level of site-and locus-filtering. Conversely, supermatrix-based analyses strongly supported the eudicots-magnoliids sister relationship when all data were included, but strongly supported the eudicots-monocots sister relationship when 54–76% of the most rapidly evolving sites were removed.

**Figure 7.**
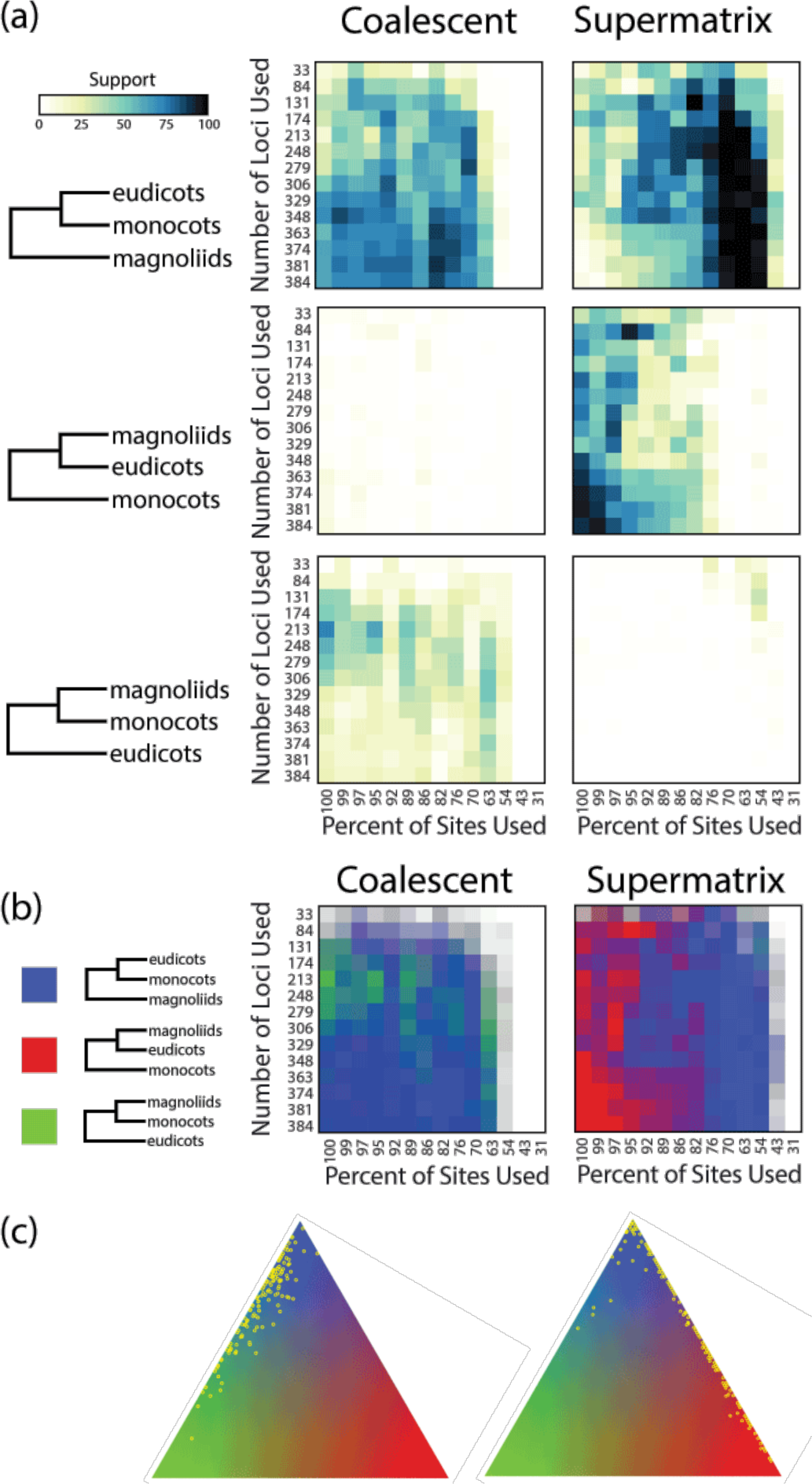
Using heat maps to compare robustness of support for key nodes across analysis frameworks. (a) Support for three alternative relationships among eudicots, monocots, and magnoliids under coalescent and supermatrix frameworks. Under the coalescent framework, a eudicot-monocot sister relationship is supported across a broad range of conditions (>63% of sites are included). Under the supermatrix approach, however, support shifts from strong support for a eudicot-magnoliid sister relationship when all sites are included to strong support for a eudicot-monocot sister relationship when 54%-76% of sites are included. (b) An additional approach for visualizing support for alternative hypotheses. In this representation, support for each of the three alternatives is indicated by a different color in the rgb (red-green-blue) color space, with the opacity of the color indicating the sum of support across the three alternatives (white indicates no support for any hypothesis). (c) A triangle representing rgb space can also be used to indicate support for alternative hypotheses. In this representation, each vertex of the triangle correponds to 100% suport for one of the three alternative hypotheses (colors have same meaning as in [b]). For each one of the data sampling schemes (pixel in [b]) a yellow point is plotted on the triangle, at the location corresponding to the relative support for the three hypotheses (a point in the middle would indicate 33% support for each of the three alternative hypotheses, whereas a point half-way along one of the edges of the triangle would indicate 50% support for each of two alternatives).

### Testing Alternative Hypotheses

Statistical tests indicated that the eudicots and monocots sister relationship was the best-supported alternative (compared to eudicots-magnoliids sister and compared to monocots-magnoliids sister) when results were averaged across all site-and locus-filtering levels. This result was obtained regardless of whether a coalescent (test statistic eudicots-monocots vs. eudicots-magnoliids = 25.09; eudicots-monocots vs. monocots-magnoliids = 52.88; p < 0.0001, both tests) or supermatrix (test statistic eudicots-monocots vs. eudicots-magnoliids = 42.63 eudicots-monocots vs. monocots-magnoliids = 24.83; p < 0.0001, both tests) approach was applied. This result may seem somewhat surprising given the inconsistent support generated by the supermatrix and coalescent approaches for this node (Fig. 7). Recall, however, that strong support for the eudicots-magnoliids sister hypothesis was only found in the narrow condition in which >95% of the sites were included in the supermatrix analysis. Support for alternative relationships of these taxa was found to be more sensitive to data filtering when the supermatrix instead of the coalescent approach was employed (test statistic = 38.69, p < 0.0001). The approximately unbiased (AU) test (Shimodaira and Hasegawa 2001) rejected the monocots-magnoliids sister hypothesis (p = 0.015), but failed to reject the eudicots-magnoliids sister hypothesis (p = 0.119).

Although estimates based on supermatrix and coalescent approaches were found to be at odds for some of the difficult nodes for some data-filtration levels, estimates based on the two approaches were quite consistent overall as long as ~150 or more loci were included. For example, analysis of the full dataset under the supermatrix and coalescent approaches yielded identical topologies within monocots and magnoliids (some topological differences were evident within eudicots, however). Interestingly, inclusion of more than 150 loci did not appear to improve the consistency of estimates between supermatrix and coalescent approaches.

## DISCUSSION

The anchored hybrid enrichment resource developed here provides systematists with the means to collect hundreds of phylogenomic loci for resolving species relationships across angiosperms. By leveraging available genomic resources we have developed an enrichment resource containing probes that represent much of the ordinal-level diversity within angiosperms, which facilitates efficient enrichment of up to 499 anchor loci. The targeted anchor loci are sufficiently conserved to allow enrichment in most groups within the clade, yet are still contain enough variation to robust resolution of the large majority of nodes at a broad range of phylogenetic scales. For instance, Léveillé-Bourret et al. (in review) demonstrate the success of the methodology in resolving the most difficult nodes of a rapid Cyperaceae radiation involving thousands of species. Our data indicate that resolution at intermediate and shallow scales is improved by including sequence data from flanking regions around the probe region because the flanks contain a greater degree of variation. Given the broad taxonomic span, this resource has the potential to unify the field of angiosperm systematics by providing a common set of nuclear loci that have a high level of phylogenetic information across evolutionary scales. In fact, the angiosperm probe v.1 kit has already been widely adopted through 21 ongoing collaborations to collect data from 87 angiosperm families for more than 2000 taxa (as of October 2016) at the Center for Anchored Phylogenomics (www.anchoredphylogeny.com).

Our results indicate that the anchor loci targeted by the angiosperm enrichment probes contain a sufficient amount of data to resolve relationships at a broad set of taxonomic scales. More than 75% of the nodes in this angiosperm-wide test were strongly supported under a broad range of conditions as long as at least 60% of the sites and 25% of the loci were included. Of the remaining nodes, only a small minority required hundreds of loci to achieve strong support. Léveillé-Bourret et al. (in review) found highly similar results in a tribal-level study, with the backbone topology stabilizing and support peaking with 100–200 loci in both concatenation and coalescent-based analyses. This pattern is not surprising given results from simulations indicating that only a few informative loci are needed to resolve easy nodes and a few hundred informative loci are sufficient to resolve more difficult nodes (Leaché and Rannala 2011). Despite the robustness of support for the majority of nodes, support values for some nodes were quite sensitive to the number of loci included, the number of sites included, and/or the analysis approach employed (supermatrix or coalescent). In most of these cases, the support peaked when an intermediate number of loci were included, suggesting that increasing the number of targeted loci beyond 499 may have a small effect on the accuracy or precision of estimates. Focusing resources on other aspects of study design, such as density of taxon sampling or increasing locus lengths, may prove to be more fruitful (Zwickl and Hillis 2002; Betancur-R. et al. 2014; Peloso et al. 2016; Simmons et al. 2016). For very shallow and/or rapidly-radiating groups, for example, robust estimates may require increased locus lengths in order to ensure well-resolved and well-supported gene trees. One approach to improve gene tree resolution for shallow-scale studies is to extend probe regions into flanks using low-coverage genome data; based on our experience, >15x coverage is typically necessary. We have had good success with this approach in orchids (Gravendeel, et al., *in prep*), lobelias (Givnish et al., *in prep*), lilies (Givnish et al., *in prep*), and goldenrods (Beck et al. *in prep*). The extended loci contain more variable regions and yield more strongly supported gene and species trees. Extending anchor loci is often preferable to targeting anonymous loci because inclusion of the anchor regions allows the data to also be useful for broad-scale meta-studies.

Despite our efforts to incorporate genomic resources from across angiosperms, enrichment efficiency could be improved further in underrepresented regions of the tree (e.g., magnoliids and basal eudicots). Fortunately, it is fairly straightforward to improve the resource by incorporating additional probes representing additional taxa as genomic resources become available. For example, as part of the orchid, lobelia, and lily projects described above, we have added probes to improve the Angiosperm v.1 kit. In each of these cases, inclusion of the additional probes substantially increased enrichment efficiency for samples from those groups (Lemmon et al., unpub. data).

The heat map approach representing robustness of phylogenetic estimates is useful in several contexts. First and foremost, it is a useful means to separate nodes with tenuous support from those with robust support. As seen with several difficult nodes in the angiosperm tree, high support values occurring under a narrow range of conditions (e.g., no site or locus filtering) has the potential to be misleading, since alternative hypotheses can be supported across a broader range of conditions. Second, the approach provides a framework for testing the robustness and consistency of alternative methods of data analysis (e.g., supermatrix and coalescent). For the monocot-eudicot-magnoliid split, for example, our results support the claim of Edwards et al. (2016) that the supermatrix approach has the tendency to produce confident yet conflicting results in different regions of parameter space, although other interpretation are possible (Simmons and Gatesy, 2016). In contrast, while the coalescent approach tends to require more data to attain high support for a hypothesis, it has a much lower tendency to produce conflicting results under different conditions. Given the potential for obtaining strong but unstable support under some conditions, we recommend utilization of the heat map approach we develop here to ensure more thorough analyses in difficult regions of the tree. Adoption of the heat-map approach may temper the claims of systematists favoring relationships that appear to be strongly supported yet are sensitive to analysis conditions or the particular choice of sites and loci to include in their dataset.

We made a substantial effort to target only low-copy loci when developing the AHE resource. Use of low-copy loci in hybrid enrichment-based phylogenomics is especially important because probes enrich not only the target regions, but also regions with moderate to high sequence similarity to the target regions. Paralogous gene regions will be enriched to a degree inversely proportional to their evolutionary divergence from the target region (Lemmon et al. 2012). In order to minimize potential error arising when enriched sequences are assessed for orthology, we estimated copy number in 45 divergent angiosperms species prior to selecting target loci for the Angiosperm v.1 design. Use of a much smaller set of taxa tends to result in an over-fitting problem wherein the taxa happen by chance to each have a single copy for some portion of loci (e.g., due to coincidental gene loss). Unsurprisingly, many of the loci previously identified as low copy based on only four taxa (*Arabidopsis, Populus, Vitis*, and *Oryza*; Duarte et al. 2010) were found to have a moderate to high number of copies in other angiosperm species (Table 3). This finding largely explains why many of the initial 959 loci identified by Duarte et al. (2010) were removed from our locus list as we refined our target set to 499 loci. The potential for over-fitting is especially likely in taxa with a history of whole genome duplications and subsequent gene loss (Glasauer and Neuhauss 2014). In teleost fishes, for example, identification of a substantial number of single-copy hybrid enrichment targets proved to be impossible despite work by Li et al. (2007), who identified 154 putatively single copy loci. In this case, Stout et al. (*in press*) circumvented this problem by developing a teleost-wide enrichment kit that targets up to six homologous copies per locus and then sorting out the gene copies into orthologs bioinformatically after sequencing. Analysis of enriched loci in teleosts and other taxa with extensive gene duplications/losses may require simultaneous estimation of speciation and duplication history (Cui et al. 2006; De Smet et al. 2013).

Many of the published studies utilizing hybrid enrichment in angiosperms have targeted the plastome, either exclusively or in combination with nuclear loci (Table 1). One challenge with simultaneously targeting the plastome and nuclear genome is overcoming the unequal coverage resulting from the large disparity in copy number between the two genomes. Some researchers desiring data from both genomes have mitigated the coverage disparity by performing two separate enrichments using kits designed separately for the two genomes (Heyduk et al. 2016; Sass et al. 2016). While this approach can be successful, it is more expensive and time consuming than AHE, and it also requires some knowledge of approximate plastid copy number. An alternative approach that has only recently been discovered is to take advantage of the fact that enrichment of nuclear targets is not perfectly efficient. Since some fraction of the sequenced library fragments are from off-target regions, it is reasonable to expect that some portion of this by-catch will be derived from the plastid genome. Indeed, one can often assemble whole plastid genomes from the enrichment by-catch (Weitemierx x et al. 2014; Stephens et al. 2015a, 2015b; Schmickl et al. 2016). Although the success of this approach depends on plastome copy number and enrichment efficiency of targeted nuclear loci, the levels of enrichment we obtained across angiosperms (2%-22% on target; Table 3) suggest that the by-catch of AHE in angiosperms is likely to contain sufficient plastome sequence reads to enable reliable reconstruction of whole plastid genomes. Note, however, that this expectation may not be met when target taxa are very closely related to one of the 25 references used to develop the probes, since enrichment efficiency in these taxa is expected to be highest. Although reconstruction of plastid genomes is not a focus of this study, the raw reads generated during enrichment of the 53 taxa used in this study could be mined for plastid genomes. For example, using a reference species from the clade as a guide, assemblies were easily constructed for the plastid genome in Cyperaceae (Buddenhagen, unpubl. data).

The diversification of angiosperm lineages continues to be debated despite the application of large genomic data sets and sophisticated phylogenetic models, leading some to wonder if we are closer to resolving Darwin’s (1903) “abominable mystery”. Our study demonstrates that despite the conflict in recent studies of deep angiosperm relationships (e.g., Duarte et al. 2010; Lee et al. 2011; Wickett et al. 2014), one hypothesis predominates (eudicots-monocots sister) over the majority of parameter space under both supermatrix and species tree methods of inference. This relationship is supported by the most recent studies focusing on the plastid genome (Moore et al. 2007; Soltis et al. 2011; Ruhfel et al. 2014) and the Angiosperm Phylogeny Group’s recent synthesis (APG IV 2016). The predominance of one hypothesis is not obvious at first glance, however, because an alternative hypothesis (i.e., eudicots-magnoliids sister) is strongly supported under a narrow but commonly chosen set of conditions (supermatrix with < 5% of sites removed). The placement of Malpighiales within the Malvids has also been debated (APG IV 2016). Despite support for alternative placement of Malpighiales by the Angiosperm Phylogeny Group, our analysis suggests robust support for Malpighiales within Malvids across a broad range of conditions. Our heat map-based analyses of robustness indicates that the differences between alternative studies of these and other deep angiosperm relationships could be reconciled through more detailed exploration of the sensitivity of each study’s findings to the particular conditions under which these studies were conducted. One especially interesting extension to the study presented herein would be to incorporate the degree of taxon sampling into the analysis of robustness, especially with respect to inclusion of the early branching taxa that were admittedly under-sampled in this study.

The taxa sampled in this study were chosen to allow the enrichment efficiency of the probe kit and the phylogenomic signal to be assessed at deep, intermediate, and shallow scales. Data from additional taxa were obtained from available genomic resources in order to increase the coverage across the angiosperm clade. The phylogeny estimated using these samples was useful for demonstrating the utility of the heat map approach at identifying robust phylogenetic estimates and for testing alternative hypotheses. Nonetheless, the taxon sampling could certainly be improved especially for basal lineages. Although the sensitivity of the phylogenetic estimates to differential taxon sampling could be assessed using the heat map approach, such an analysis is outside the scope of this paper, and so we leave this effort for future studies.

## CONCLUDING REMARKS

The anchored enrichment resource we developed provides an efficient way of obtaining a sufficient amount of phylogenomic data for deep, intermediate, and shallow angiosperm studies. Although most angiosperm relationships could be robustly resolved, support for a small minority of relationships varied substantially depending on the particular subset of sites and loci analyzed, suggesting that the sensitivity approach we developed here may helpful for avoiding false confidence in hypotheses that supported under only a narrow set of conditions. Use of this holistic approach may enable systematists to reconcile differences across studies as they work towards a unified understanding the history of angiosperm diversification.

## ACKNOWLEDGEMENTS

We are grateful to Alyssa Bigelow, Kirby Birch, Ameer Jalal, and Sean Holland at FSU’s Center for Anchored Phylogenomics for assistance with molecular data collection and bioinformatics analysis. We are also grateful to Roger Mercer and Yanming Yang at FSU’s College of Medicine for sequencing services. This research was partially funded by NSF DDIG (DEB-1311150) to CEB and by a FSU Planning Grant awarded to ARM. EML acknowledges support from NSF DEB-1120516. ARL and EML were supported by NSF IIP-1313554. CP would like to acknowledge the generous support of the Hermon Slade Foundation, Emily Holmes Memorial Scholarship; Hansjörg Eichler Scientific Research Fund; Systematics Research Fund, without which research material used in this publication would not have been collected. NM was supported by NSF DEB-1046328. SW thanks the TU Dresden and especially Christoph Neinhuis for continuous support. AM was supported by NSF DEB-1252901 to EJE. PSS is grateful for support from iDigBio (NSF grant EF-1115210) and a Dimensions of Biodiversity grant (DEB-1442280).

## LITERATURE CITED

APG IV 2016. An update of the Angiosperm Phylogeny Group classification for the orders and families of flowering plants: APG IV. Bot. J. Linn. Soc. 181:1–20.

Bakker F.T. 2015. DNA sequences from plant herbarium tissue. In: Appelhans M.S., Hörandl E., editors. Next-Generation Sequencing in Plant Systematics. International Association for Plant Taxonomy.

Beck J.B., Semple J.C. 2015. Next-generation sampling: Pairing genomics with herbarium specimens provides species-level signal in *Solidago* (Asteraceae). Appl. Plant Sci. 3:apps.1500014.

Betancur-R. R., Naylor G.J.P., Ortí G. 2014. Conserved Genes, Sampling Error, and Phylogenomic Inference. Syst. Biol. 63:257–262.

Bi K., Linderoth T., Vanderpool D., Good J.M., Nielsen R., Moritz C. 2013. Unlocking the vault: next-generation museum population genomics. Mol. Ecol. 22:6018–6032.

Bogdanowicz D., Giaro K., Wróbel B. 2012. TreeCmp: Comparison of trees in polynomial time. Evol. Bioinform. 8:475–487.

Brandley M.C., Bragg J.G., Singhal S., Chapple D.G., Jennings C.K., Lemmon A.R., Lemmon E.Moriarty, Thompson M.B., Moritz C. 2015. Evaluating the performance of anchored hybrid enrichment at the tips of the tree of life: a phylogenetic analysis of Australian *Eugongylus* group scincid lizards. BMC Evol. Biol. 15:1–14.

Breinholt J.W., Kawahara A.Y. 2013. Phylotranscriptomics: Saturated Third Codon Positions Radically Influence the Estimation of Trees Based on Next-Gen Data. Genome Biol. Evol. 5:2082–2092.

Breinholt, J.W., Lemmon A.R., Lemmon E.M., Xiao L., and Kawahara A.Y. Anchored hybrid enrichment in Lepidoptera: Leveraging genomic data for studies on the megadiverse butterflies and moths. Syst. Biol. *Accepted.*

Brown J.M., Lemmon A.R. 2007. The importance of data partitioning and the utility of Bayes factors in Bayesian phylogenetics. Syst. Biol. 56:643–655.

Bryson Jr R.W., Faircloth B.C., Tsai W.L., McCormack J.E., Klicka J. 2016. Target enrichment of thousands of ultraconserved elements sheds new light on early relationships within New World sparrows (Aves: Passerellidae). The Auk. 133:451–458.

Burleigh J.G., Bansal M.S., Eulenstein O., Hartmann S., Wehe A., Vision T.J. 2011. Genome-Scale Phylogenetics: Inferring the Plant Tree of Life from 18,896 Gene Trees. Syst. Biol. 60:117–125.

Castresana J. 2000. Selection of Conserved Blocks from Multiple Alignments for Their Use in Phylogenetic Analysis. Mol. Biol. Evol. 17:540–552.

Chase M.W., Soltis D.E., Olmstead R.G., Morgan D., Les D.H., Mishler B.D., Duvall M.R., Price R.A., Hills H.G., Qiu Y.-L., Kron K.A., Rettig J.H., Conti E., Palmer J.D., Manhart J.R., Sytsma K.J., Michaels H.J., Kress W.J., Karol K.G., Clark W.D., Hedren M., Gaut B.S., Jansen R.K., Kim K-J., Wimpee C.F., Smith J.F., Furnier G.R., Strauss S.H., Xiang Q-Y, Plunkett G.M., Soltis P.S., Swensen S.M., Williams S.E., Gadek P.A., Quinn C.J., Eguiarte L.E., Golenberg E., Learn Jr. G.H., Graham S.W., Barrett S.C.H., Dayanandan S., Albert V.A. 1993. Phylogenetics of seed plants: an analysis of nucleotide sequences from the plastid gene rbcL. Ann. Mo. Bot. Gard. 80:528–580.

Chen F., Mackey A.J., Vermunt J.K., Roos D.S. 2007. Assessing performance of orthology detection strategies applied to eukaryotic genomes. PloS One. 2:e383.

Costa C.M., Roberts R.P. 2014. Techniques for improving the quality and quantity of DNA extracted from herbarium specimens. Phytoneuron. 48:1–8.

Crawford N.G., Faircloth B.C., McCormack J.E., Brumfield R.T., Winker K., Glenn T.C. 2012. More than 1000 ultraconserved elements provide evidence that turtles are the sister group of archosaurs. Biol. Lett. 8:783–786.

Crawford N.G., Parham J.F., Sellas A.B., Faircloth B.C., Glenn T.C., Papenfuss T.J., Henderson J.B., Hansen M.H., Simison W.B. 2015. A phylogenomic analysis of turtles. Mol. Phylogenet. Evol. 83:250–257.

Critchlow D.E., Pearl D.K., Qian C. 1996. The triples distance for rooted bifurcating phylogenetic trees. Syst. Biol. 45:323–334.

Cronn R., Knaus B.J., Liston A., Maughan P.J., Parks M., Syring J.V., Udall J. 2012. Targeted enrichment strategies for next-generation plant biology. Am. J. Bot. 99:291–311.

Cui L., Wall P.K., Leebens-Mack J.H., Lindsay B.G., Soltis D.E., Doyle J.J., Soltis P.S., Carlson J.E., Arumuganathan K., Barakat A. 2006. Widespread genome duplications throughout the history of flowering plants. Genome Res. 16:738–749.

Darriba D., Posada D. 2015. The impact of partitioning on phylogenomic accuracy. bioRxiv doi: http://dx.doi.org/10.1101/023978.

Darwin C., Darwin F., and Seward A.C. 1903. More letters of Charles Darwin: a record of his work in a series of hitherto unpublished letters. D. Appleton and Co., New York.

Davis C.C., Xi Z., Mathews S. 2014. Plastid phylogenomics and green plant phylogeny: almost full circle but not quite there. BMC Biol. 12:1–4.

De Smet R., Adams K.L., Vandepoele K., Van Montagu M.C.E., Maere S., Van de Peer Y. 2013. Convergent gene loss following gene and genome duplications creates single-copy families in flowering plants. Proc. Natl. Acad. Sci. 110:2898–2903.

Duarte J.M., Wall P.K., Edger P.P., Landherr L.L., Ma H., Pires J.C., Leebens-Mack J. 2010. Identification of shared single copy nuclear genes in Arabidopsis, Populus, Vitis and Oryza and their phylogenetic utility across various taxonomic levels. BMC Evol. Biol. 10:61.

Edwards S.V. 2009. Is a new and general theory of molecular systematics emerging? Evolution. 63:1–19.

Edwards S.V. 2016. Phylogenomic subsampling: a brief review. Zoologica Scripta 45: 63–74.

Edwards S.V., Xi Z., Janke A., Faircloth B.C., McCormack J.E., Glenn T.C., Zhong B., Wu S., Lemmon E.M., Lemmon A.R. 2016. Implementing and testing the multispecies coalescent model: a valuable paradigm for phylogenomics. Mol. Phylogenet. Evol. 94:447–462.

Eytan R.I., Evans B.R., Dornburg A., Lemmon A.R., Lemmon E.M., Wainwright P.C., Near T.J. 2015. Are 100 enough? Inferring acanthomorph teleost phylogeny using Anchored Hybrid Enrichment. BMC Evol. Biol. 15:1.

Faircloth B.C., McCormack J.E., Crawford N.G., Harvey M.G., Brumfield R.T., Glenn T.C. 2012. Ultraconserved elements anchor thousands of genetic markers spanning multiple evolutionary timescales. Syst. Biol. 61:717–726.

Frandsen P.B., Calcott B., Mayer C., Lanfear R. 2015. Automatic selection of partitioning schemes for phylogenetic analyses using iterative k-means clustering of site rates. BMC Evol. Biol. 15:1–17.

Freeman J.L., Tamaoki M., Stushnoff C., Quinn C.F., Cappa J.J., Devonshire J., Fakra S.C., Marcus M.A., McGrath S.P., Van Hoewyk D., Pilon-Smits E.A.H. 2010. Molecular mechanisms of selenium tolerance and hyperaccumulation in *Stanleya pinnata*. Plant Physiol. 153:1630–1652.

Glasauer S.M., Neuhauss S.C. 2014. Whole-genome duplication in teleost fishes and its evolutionary consequences. Mol. Genet. Genomics. 289:1045–1060.

Griffin P., Robin C., Hoffmann A. 2011. A next-generation sequencing method for overcoming the multiple gene copy problem in polyploid phylogenetics, applied to *Poa* grasses. BMC Biol. 9:1–19.

Grover C.E., Gallagher J.P., Jareczek J.J., Page J.T., Udall J.A., Gore M.A., Wendel J.F. 2015. Re-evaluating the phylogeny of allopolyploid *Gossypium* L. Molecular Phylogenetics and Evolution. 92:45–52.

Hamilton C.A., Lemmon A.R., Lemmon E. Moriarty, Bond J.E. 2016. Expanding anchored hybrid enrichment to resolve both deep and shallow relationships within the spider tree of life. BMC Evolutionary Biology 16:212.

Heyduk K., Trapnell D.W., Barrett C.F., Leebens-Mack J. 2016. Phylogenomic analyses of species relationships in the genus *Sabal* (Arecaceae) using targeted sequence capture. Biol. J. Linn. Soc. 117:106–120.

Hillis D.M., Huelsenbeck J.P., Cunningham C.W. 1994. Application and accuracy of molecular phylogenies. Science 264:671–677.

Hoff M., Orf S., Riehm B., Darriba D., Stamatakis A. 2016. Does the choice of nucleotide substitution models matter topologically? BMC Bioinformatics 17:1–13.

Hosner P.A., Faircloth B.C., Glenn T.C., Braun E.L., Kimball R.T. 2015. Avoiding missing data biases in phylogenomic inference: an empirical study in the landfowl (Aves: Galliformes). Mol. Biol. Evol. 33:1110–1125.

Jiao Y., Wickett N.J., Ayyampalayam S., Chanderbali A.S., Landherr L., Ralph P.E., Tomsho L.P., Hu Y., Liang H., Soltis P.S., Soltis D.E., Clifton S.W., Schlarbaum S.E., Schuster S.C., Ma H., Leebens-Mack J., dePamphilis C.W. 2011. Ancestral polyploidy in seed plants and angiosperms. Nature. 473:97–100.

Katoh K., Standley D.M. 2013. MAFFT multiple sequence alignment software version 7: improvements in performance and usability. Mol. Biol. Evol. 30:772–780.

Kearse M., Moir R., Wilson A., Stones-Havas S., Cheung M., Sturrock S., Buxton S., Cooper A., Markowitz S., Duran C. 2012. Geneious Basic: an integrated and extendable desktop software platform for the organization and analysis of sequence data. Bioinformatics. 28:1647–1649.

Kluge A.G. 1989. A concern for evidence and a phylogenetic hypothesis of relationships among *Epicrates* (Boidae, Serpentes). Syst. Biol. 38:7–25.

Lanfear R., Calcott B., Ho S.Y., Guindon S. 2012. PartitionFinder: combined selection of partitioning schemes and substitution models for phylogenetic analyses. Mol. Biol. Evol. 29:1695–1701.

Leaché A.D., Rannala B. 2011. The accuracy of species tree estimation under simulation: a comparison of methods. Syst. Biol. 60:126–137.

Lee E.K., Cibrian-Jaramillo A., Kolokotronis S.-O., Katari M.S., Stamatakis A., Ott M., Chiu J.C., Little D.P., Stevenson D.W., McCombie W.R. 2011a. A functional phylogenomic view of the seed plants. PLoS Genet. 7:e1002411.

Lemmon A.R., Brown, J.M., Stanger-Hall, K., Lemmon, E. Moriarty. 2009. The effect of ambiguous data on phylogenetic estimates obtained by maximum likelihood and Bayesian inference. Syst. Biol. 58:130–145.

Lemmon A.R., Emme S.A., Lemmon E. Moriarty 2012. Anchored Hybrid Enrichment for Massively High-Throughput Phylogenomics. Syst. Biol. 61:727–744.

Lemmon A.R., Moriarty E.C. 2004. The importance of proper model assumption in Bayesian phylogenetics. Syst. Biol. 53:265–277.

Lemmon E.M., Lemmon A.R. 2013. High-Throughput Genomic Data in Systematics and Phylogenetics. Annu. Rev. Ecol. Evol. Syst. 44:99–121.

Lemon J. 2006. Plotrix: a package in the red light district of R. R-News. 6:8–12.

Léveillé-Bourret E., Starr J.R., Ford B.A., Lemmon E.M., Lemmon A.R. *In review*. Anchored phylogenomics for angiosperms II: Resolving rapid generic and tribal-level radiations. Syst. Biol.

Li C., Ortí G., Zhang G., Lu G. 2007. A practical approach to phylogenomics: the phylogeny of ray-finned fish (Actinopterygii) as a case study. BMC Evol. Biol. 7:44–44.

Magallón S., Hilu K.W., Quandt D. 2013. Land plant evolutionary timeline: Gene effects are secondary to fossil constraints in relaxed clock estimation of age and substitution rates. Am. J. Bot. 100: 556–573.

Mamanova L., Coffey A.J., Scott C.E., Kozarewa I., Turner E.H., Kumar A., Howard E., Shendure J., Turner D.J. 2010. Target-enrichment strategies for next-generation sequencing. Nat. Methods. 7:111–118.

Mandel J.R., Dikow R.B., Funk V.A., Masalia R.R., Staton S.E., Kozik A., Michelmore R.W., Rieseberg L.H., Burke J.M. 2014. A Target Enrichment Method for Gathering Phylogenetic Information from Hundreds of Loci: An Example from the Compositae. Appl. Plant Sci. 2:1300085.

McCormack J.E., Faircloth B.C., Crawford N.G., Gowaty P.A., Brumfield R.T., Glenn T.C. 2012. Ultraconserved elements are novel phylogenomic markers that resolve placental mammal phylogeny when combined with species-tree analysis. Genome Res. 22:746–754.

Meiklejohn K.A., Faircloth B.C., Glenn T.C., Kimball R.T., Braun E.L. 2016. Analysis of a Rapid Evolutionary Radiation Using Ultraconserved Elements: Evidence for a Bias in Some Multispecies Coalescent Methods. Syst. Biol. 65:612–627.

Meredith R.W., Janecka J.E., Gatesy J., Ryder O.A., Fisher C.A., Teeling E.C., Goodbla A., Eizirik E., Simao T.L.L., Stadler T., Rabosky D.L., Honeycutt R.L., Flynn J.J., Ingram C.M., Steiner C., Williams T.L., Robinson T.J., Burk-Herrick A., Westerman M., Ayoub N.A., Springer M.S., Murphy W.J. 2011. Impacts of the Cretaceous terrestrial revolution and KPg extinction on mammal diversification. Science 334:521–524.

Meyer M., Kircher M. 2010. Illumina sequencing library preparation for highly multiplexed target capture and sequencing. Cold Spring Harb. Protoc. 2010; doi:10.1101/pdb.prot5448.

Mirarab S., Warnow T. 2015. ASTRAL-II: coalescent-based species tree estimation with many hundreds of taxa and thousands of genes. Bioinformatics. 31:i44–i52.

Moore M.J., Bell C.D., Soltis P.S., Soltis D.E. 2007. Using plastid genome-scale data to resolve enigmatic relationships among basal angiosperms. Proc. Natl. Acad. Sci. 104:19363–19368.

Morales H.E., Pavlova A., Sunnucks P., Major R., Amos N., Joseph L., Lemmon A.R., Endler J.A., Delhey K. Neutral and selective drivers of colour evolution in a widespread Australian passerine. J. Biogeogr.; http://hdl.handle.net/11858/00-001M-0000-002B-11FF-D *In review*

Morrison D. 2011. Introduction to Phylogenetic Networks. RJR Publications. Uppsala, Sweden.

Paradis E., Claude J., Strimmer K. 2004. APE: analyses of phylogenetics and evolution in R language. Bioinformatics. 20:289–290.

Parks M., Cronn R., Liston A. 2012. Separating the wheat from the chaff: mitigating the effects of noise in a plastome phylogenomic data set from *Pinus* L. (Pinaceae). BMC Evol. Biol. 12:1–17.

Peloso P.L.V., Frost D.R., Richards S.J., Rodrigues M.T., Donnellan S., Matsui M., Raxworthy C.J., Biju S.D., Lemmon E.M., Lemmon A.R., Wheeler W.C. 2016. The impact of anchored phylogenomics and taxon sampling on phylogenetic inference in narrow-mouthed frogs (Anura, Microhylidae). Cladistics 32:113–140.

Pimm S.L., Joppa L.N. 2015. How many plant species are there, where are they, and at what rate are they going extinct? Ann. Mo. Bot. Gard. 100:170–176.

Posada D., Crandall K.A. 1998. Modeltest: testing the model of DNA substitution. Bioinformatics 14:817–818.

Prum R.O., Berv J.S., Dornburg A., Field D.J., Townsend J.P., Lemmon E.M., Lemmon A.R. 2015. A comprehensive phylogeny of birds (Aves) using targeted next-generation DNA sequencing. Nature. 526:569–573.

Pyron R.A., Hendry C.R., Chou V.M., Lemmon E.M., Lemmon A.R., Burbrink F.T. 2014. Effectiveness of phylogenomic data and coalescent species-tree methods for resolving difficult nodes in the phylogeny of advanced snakes (Serpentes: Caenophidia). Mol. Phylogenet. Evol. 81:221–231.

de Queiroz A., Gatesy J. 2007. The supermatrix approach to systematics. Trends Ecol. Evol. 22:34–41.

Reneker J., Lyons E., Conant G.C., Pires J.C., Freeling M., Shyu C.-R., Korkin D. 2012. Long identical multispecies elements in plant and animal genomes. Proc. Natl. Acad. Sci. 109:E1183–E1191.

Ripplinger J., Sullivan J. 2008. Does choice in model selection affect maximum likelihood analysis? Syst. Biol. 57:76–85.

Rokyta D.R., Lemmon A.R., Margres M.J., Aronow K. 2012. The venom-gland transcriptome of the eastern diamondback rattlesnake (*Crotalus adamanteus*). BMC Genomics. 13:312.

Roure B., Baurain D., Philippe H. 2013. Impact of missing data on phylogenies inferred from empirical phylogenomic data sets. Mol. Biol. Evol. 30:197–214.

Ruane S., Raxworthy C.J., Lemmon A.R., Lemmon E.M., Burbrink F.T. 2015. Comparing species tree estimation with large anchored phylogenomic and small Sanger-sequenced molecular datasets: an empirical study on Malagasy pseudoxyrhophiine snakes. BMC Evol. Biol. 15:1.

Ruhfel B.R., Gitzendanner M.A., Soltis P.S., Soltis D.E., Burleigh J.G. 2014. From algae to angiosperms-inferring the phylogeny of green plants (Viridiplantae) from 360 plastid genomes. BMC Evol. Biol. 14:23.

Sass C., Iles W.J.D., Barrett C.F., Smith S.Y., Specht C.D. 2016. Revisiting the Zingiberales: using multiplexed exon capture to resolve ancient and recent phylogenetic splits in a charismatic plant lineage. Peerj. 4:e1584.

Schmickl R., Liston A., Zeisek V., Oberlander K., Weitemier K., Straub S.C., Cronn R.C., Dreyer L.L., Suda J. 2016. Phylogenetic marker development for target enrichment from transcriptome and genome skim data: the pipeline and its application in southern African *Oxalis* (Oxalidaceae). Mol. Ecol. Resour. 16:1124–1135.

Shimodaira H., Hasegawa M. 2001. CONSEL: for assessing the confidence of phylogenetic tree selection. Bioinformatics. 17:1246–1247.

Shimodaira, H. 2002. An approximately unbiased test of phylogenetic tree selection. Syst. Biol. 51: 492–508.

Simmons, M. P., J. Gatesy. 2015. Coalescence vs. concatenation: sophisticated analyses vs. first principles applied to rooting the angiosperms. Mol. Phylogenet. Evol. 91: 98–122.

Simmons, M. P., J. Gatesy. 2016. Biases of tree-independent-character-subsampling methods. Mol. Phylogenet. Evol. 100: 424–443.

Simmons M.P., Sloan D.B., Gatesy J. 2016. The effects of subsampling gene trees on coalescent methods applied to ancient divergences. Mol. Phylogenet. Evol. 97:76–89.

Song S., Liu L., Edwards S.V., Wu S., 2012. Resolving conflict in eutherian mammal phylogeny using phylogenomics and the multispecies coalescent model. Proc. Natl. Acad. Sci. USA 109:14942–14947.

Springer M. S., J. Gatesy. 2016. The gene tree delusion. Mol. Phylogenet. Evol. 94:1–33.

Soltis D.E., Smith S.A., Cellinese N., Wurdack K.J., Tank D.C., Brockington S.F., Refulio-Rodriguez N.F., Walker J.B., Moore M.J., Carlsward B.S. 2011. Angiosperm phylogeny: 17 genes, 640 taxa. Am. J. Bot. 98:704–730.

de Sousa F., Bertand Y.J.K., Nylinder S., Oxelman B., Eriksson J.S., Pfeil B.E. 2014. Phylogenetic properties of 50 nuclear loci in *Medicago* (Leguminosae) generated using multiplexed sequence capture and next-generation sequencing. PLoS ONE 9:e109704. doi: 10.1371/journal.pone.0109704.

Stamatakis A. 2014. RAxML version 8: a tool for phylogenetic analysis and post-analysis of large phylogenies. Bioinformatics. 30:1312–1313.

Stephens J.D., Rogers W.L., Heyduk K., Cruse-Sanders J.M., Determann R.O., Glenn T.C., Malmberg R.L. 2015a. Resolving phylogenetic relationships of the recently radiated carnivorous plant genus *Sarracenia* using target enrichment. Mol. Phylogenet. Evol. 85:76–87.

Stephens J.D., Rogers W.L., Mason C.M., Donovan L.A., Malmberg R.L. 2015b. Species tree estimation of diploid *Helianthus* (Asteraceae) using target enrichment. Am. J. Bot. 102:910–920.

Stout, C.C., Tan M., Lemmon A.R., Lemmon E.M., Armbruster J.W. Resolving Cypriniformes Relationships Using An Anchored Enrichment Approach. BMC Evol. Biol. *In Review.*

Stull G.W., Moore M.J., Mandala V.S., Douglas N.A., Kates H.-R., Qi X., Brockington S.F., Soltis P.S., Soltis D.E., Gitzendanner M.A. 2013. A targeted enrichment strategy for massively parallel sequencing of angiosperm plastid genomes. Appl. Plant Sci. 1:1200497.

Sun Y., Moore M.J., Zhang S., Soltis P.S., Soltis D.E., Zhao T., Meng A., Li X., Li J., Wang H. 2016. Phylogenomic and structural analyses of 18 complete plastomes across nearly all families of early-diverging eudicots, including an angiosperm-wide analysis of IR gene content evolution. Mol. Phylogenet. Evol. 96:93–101.

Syring J., Cronn R., Tennessen J.A., Jennings T.N., Scelfo-Dalbey C., Wegrzyn J. 2016. Targeted capture sequencing in whitebark pine reveals range-wide demographic and adaptive patterns despite challenges of a large, repetitive genome. Frontiers in Plant Science. 7:484.

Talavera G., Castresana J. 2007. Improvement of phylogenies after removing divergent and ambiguously aligned blocks from protein sequence alignments. Syst. Biol. 56:564–577.

Berardini T.Z., Reiser L, Li D., Mezheritsky Y., Muller R., Strait E., Huala E. 2015. The Arabidopsis Information Resource: Making and mining the “gold standard” annotated reference plant genome. Genesis. 53:474–485.

Terry N., Zayed A.M., de Souza M.P., Tarun A.S. 2000. Selenium in higher plants. Ann. Rev. Plant Physiol. Plant Mol. Biol. 51:401–432.

Townsend, T.M., Mulcahy, D.G., Noonan, B.P., Sites, J.W., Kuczynski, C.A., Wiens, J.J., Reeder, T.W., 2011. Phylogeny of iguanian lizards inferred from 29 nuclear loci, and a comparison of concatenated and species-tree approaches for an ancient, rapid radiation. Mol. Phylogenet. Evol. 61:363–380.

Tucker D.B., Colli G.R., Giugliano L.G., Hedges S.B., Hendry C.R., Lemmon E. Moriarty, Lemmon A.R., Pyron R.A. 2016. Methodological congruence in phylogenomic analyses with morphological support for teiid lizards (Sauria: Teiidae). Mol. Phylogenet. Evol. 103:75–84.

Weitemier K., Straub S.C., Cronn R.C., Fishbein M., Schmickl R., McDonnell A., Liston A. 2014. Hyb-Seq: Combining target enrichment and genome skimming for plant phylogenomics. Appl. Plant Sci. 2:1400042.

Wicke S., Schneeweiss G.M. 2015. Next-generation organellar genomics: Potentials and pitfalls of high-throughput technologies for molecular evolutionary studies and plant systematics. In: Horandl E., Appelhans M.S., editors. Next Generation Sequencing in Plant Systematics. Koenigstein: A R G Gantner Verlag K G. Pp. 9–50.

Wickett N.J., Mirarab S., Nguyen N., Warnow T., Carpenter E., Matasci N., Ayyampalayam S., Barker M.S., Burleigh J.G., Gitzendanner M.A., Ruhfel B.R., Wafula E., Der J.P., Graham S.W., Mathews S., Melkonian M., Soltis D.E., Soltis P.S., Miles N.W., Rothfels C.J., Pokorny L., Shaw A.J., DeGironimo L., Stevenson D.W., Surek B., Villarreal J.C., Roure B., Philippe H., dePamphilis C.W., Chen T., Deyholos M.K., Baucom R.S., Kutchan T.M., Augustin M.M., Wang J., Zhang Y., Tian Z., Yan Z., Wu X., Sun X., Wong G.K.-S., Leebens-Mack J. 2014. Phylotranscriptomic analysis of the origin and early diversification of land plants. Proc. Natl. Acad. Sci. 111:E4859–E4868.

Xia X., Lemey P. 2009. Assessing substitution saturation with DAMBE. In: Lemey P., Salemi M., and Vandamme A.-M., editors. The Phylogenetic Handbook: a Practical Approach to Phylogenetic Analysis and Hypothesis Testing. Cambridge University Press, UK. Pp. 611–626.

Yang Z. 1994. Maximum likelihood phylogenetic estimation from DNA sequences with variable rates over sites: approximate methods. J. Mol. Evol. 39:306–314.

Young A.D., Lemmon A.R., Skevington J.H., Mengual X., Ståhls G., Reemer M., Jordaens K., Kelso S., Lemmon E. Moriarty, Hauser M. 2016. Anchored enrichment dataset for true flies (order Diptera) reveals insights into the phylogeny of flower flies (family Syrphidae). BMC Evol. Biol. 16:143.

Zeng L., Zhang Q., Sun R., Kong H., Zhang N., Ma H. 2014. Resolution of deep angiosperm phylogeny using conserved nuclear genes and estimates of early divergence times. Nat Commun. 5:4956.

Zimmer E.A., Wen J. 2012. Using nuclear gene data for plant phylogenetics: Progress and prospects. Mol. Phylogenet. Evol. 65:774–785.

Zwickl D.J., Hillis D.M. 2002. Increased taxon sampling greatly reduces phylogenetic error. Syst. Biol. 51:588–598.

